# Lipid Droplet Remodeling Safeguards Redox Balance through the DGAT1–PANK2–NRF2 Axis in Mammalian Oocytes and Drives Age-Associated Decline

**DOI:** 10.1101/2025.11.28.691261

**Authors:** Pravin Birajdar, Akshay Kumar, Anjali Kumari, Aradhana Mohanty, Mohd Athar, Ajith Kumar, Kiran Kumar. P, Abhilasha. S, Sahina Sabnam, Silambresan. Y, Ankita Verma, Rajendar. M, H.B.D. Prasada Rao

## Abstract

How lipid droplets (LDs) buffer metabolic stress and redox imbalance in aging oocytes remains poorly understood. Here, we identify de novo LD remodeling as a metabolic capacitor that couples lipid storage to mitochondrial fitness and oxidative resilience in mammalian oocytes. Live imaging revealed pronounced LD dynamics, with LD number peaking at metaphase I and declining by metaphase II, while LD area shifted inversely. Despite stable triacylglyceride and free fatty acid pools, β-oxidation increased sharply, indicating elevated lipid turnover during meiotic progression. Spatial mapping and fatty-acid tracing demonstrated that newly synthesized lipids are actively incorporated into LDs, which arise primarily from the endoplasmic reticulum and engage with lysosomes and mitochondria. Acute inhibition of DGAT1, the rate-limiting enzyme of LD biogenesis, disrupted meiotic maturation and triggered oxidative stress, mitochondrial aggregation, and ultrastructural damage. Proteomic profiling revealed robust PANK2 upregulation and suppression of NRF2-linked antioxidant pathways. Mechanistic analyses showed that β-oxidation blockade, PANK2 inhibition, antioxidant supplementation, or NRF2 activation each partially rescued DGAT1-dependent defects, and genetic validation in NRF2-null oocytes confirmed pathway dependence.

Notably, aged oocytes exhibited reduced de novo LD biogenesis and impaired DGAT1–ER organization despite increased LD accumulation, resulting in smaller, metabolically inert droplets and a mismatch between lipid formation and utilization. Inhibiting PANK2 alleviated oxidative stress in aged oocytes, further implicating the DGAT1–PANK2–NRF2 axis in redox control and oocyte quality. Together, these findings establish LD biogenesis as a core metabolic capacitor safeguarding mitochondrial and organelle integrity during meiosis and reveal dysfunction of the DGAT1–PANK2–NRF2 axis as a mechanistic driver of reproductive aging.

## Introduction

Mammalian oocyte meiotic maturation is a highly regulated developmental process where primary oocytes, initially arrested in prophase I of meiosis, resume progression to become mature, fertilization competent eggs ^1, 2^. This process involves critical events such as germinal vesicle breakdown (GVBD), chromosome condensation, spindle formation, and homologous chromosome segregation during meiosis I, followed by a transient arrest at metaphase II until fertilization ^3, 4^. Successful maturation requires substantial energy to support cellular activities such as protein synthesis, metabolic reprogramming, calcium signaling, RNA and protein storage, and cellular remodeling^5–7^. Proper regulation of these energy dependent processes is essential for oocyte maturation and subsequent fertilization. A delicate balance of lipid and sugar metabolism ensures a sustained energy supply, supporting both immediate and long-term energy needs throughout maturation^6, 8–10^.

Lipids, predominantly stored as triglycerides within cytoplasmic lipid droplets (LDs), play a pivotal role in oocyte maturation^10–13^. During this process, lipid reserves are mobilized through lipolysis, releasing free fatty acids that are transported to mitochondria for β-oxidation ^9, 14, 15^. This metabolic pathway produces acetyl-CoA, which enters the citric acid cycle to generate ATP, an essential energy source for sustaining meiotic arrest, spindle assembly, and the progression of meiosis^16–19^. Beyond their role in energy production, lipids are fundamental for membrane biogenesis, a process critical for the cytoskeletal remodeling and organelle reorganization that accompany oocyte maturation ^10, 20–22^. Recent studies have highlighted the importance of fatty acid oxidation in the early stages of maturation, ensuring a steady supply of ATP to support protein synthesis, spindle dynamics, and other energy-intensive cellular functions ^8, 14, 23^. Moreover, lipid-derived acetyl-CoA serves as a precursor for the synthesis of signaling molecules and metabolic cofactors that regulate meiotic progression^8, 22, 24, 25^.

While lipid metabolism is well recognized as a key component of oocyte maturation, the specific contributions of LDs to energy homeostasis and redox regulation remain incompletely understood. Emerging evidence suggests that LDs not only function as energy reservoirs but also participate in mitigating oxidative stress and maintaining cellular homeostasis^26–28^. LD formation begins during early folliculogenesis, around postnatal days 11 to 12 in mice, and is closely associated with the endoplasmic reticulum (ER)^29^. Tracing studies have demonstrated that oocytes rapidly take up exogenous fatty acids, a process modulated by interactions with surrounding granulosa cells^29^. However, the precise mechanisms by which LDs influence oocyte meiotic maturation and developmental competence remain an active area of investigation.

LD formation and regulation are governed by enzymes such as diacylglycerol O-acyltransferase 1 (DGAT1) and DGAT2 ^30–33^. DGAT1, located primarily in the ER, plays a critical role in storing triglycerides, thereby maintaining energy balance and preventing lipotoxicity^31, 34^. DGAT2, localized at the ER-LD junctions, facilitates the transfer of newly synthesized TAGs into lipid droplets, a process that is particularly relevant in tissues with high lipid turnover^35–37^. The regulation of lipid droplet formation by DGAT enzymes in regulating metabolism and redox balance highlights the complex interplay between lipid metabolism and antioxidant defense mechanisms in oocytes.

Oxidative stress, a byproduct of increased metabolic activity, is another key factor affecting oocyte competence^38–40^. ^41^)^39, 42^. While low levels of ROS are necessary for normal cellular signaling, excessive ROS accumulation can damage cellular components, including lipids, proteins, and DNA, impairing oocyte function^43, 44^. Moreover, advancing maternal age is associated with a decline in the oocyte’s antioxidant capacity, leading to greater ROS accumulation, reduced oocyte quality, and diminished ovarian reserve^45–48^. NRF2, a key regulator of redox homeostasis, plays a critical role in protecting oocytes from oxidative stress, a common consequence of mitochondrial dysfunction and lipid metabolism disturbances^49, 50^. ^51, 52^. This regulation is particularly important in oocytes, which are energy-demanding cells vulnerable to ROS accumulation. Recent studies have linked NRF2 signaling to lipid droplet formation, underscoring its role in coordinating lipid metabolism and maintaining redox balance during oocyte maturation ^53, 54^. Lipid droplets not only serve as energy reservoirs but also modulate oxidative stress, and their formation is tightly integrated with metabolic reprogramming^26^. This intricate balance of energy and oxidative stress is essential for maintaining oocyte quality and supporting reproductive success ^55, 56^. Thus, this study aims to explore the relationship between lipid droplet formation, energy homeostasis, and NRF2-mediated redox regulation, offering insights into the mechanisms that maintain oocyte quality and promote successful reproduction.

## Results

### Dynamic de novo lipid droplet remodeling buffers energy demand during meiotic maturation

To investigate how lipid droplets (LDs) contribute to metabolic remodeling during mammalian oocyte meiotic maturation, we visualized their dynamics using Nile Red (neutral lipids) and BODIPY (LD marker) in mouse and goat oocytes (Fig. 1A, E; Fig. S1A, C, E). LDs were tracked across germinal vesicle (GV), metaphase I (MI), and metaphase II-like (MII; anaphase/telophase I) stages (Fig. 1B, F; Fig. S1B, D, F). Quantification revealed striking dynamics: LD number rose significantly from GV to MI, then returned to near-GV levels by MII, while average LD area displayed the inverse trend (Fig. 1C–H). These opposing shifts indicate a continuous cycle of de novo LD formation and remodeling, consistent with LDs acting as a transient, flexible energy reservoir during meiosis.

**Fig 1:**
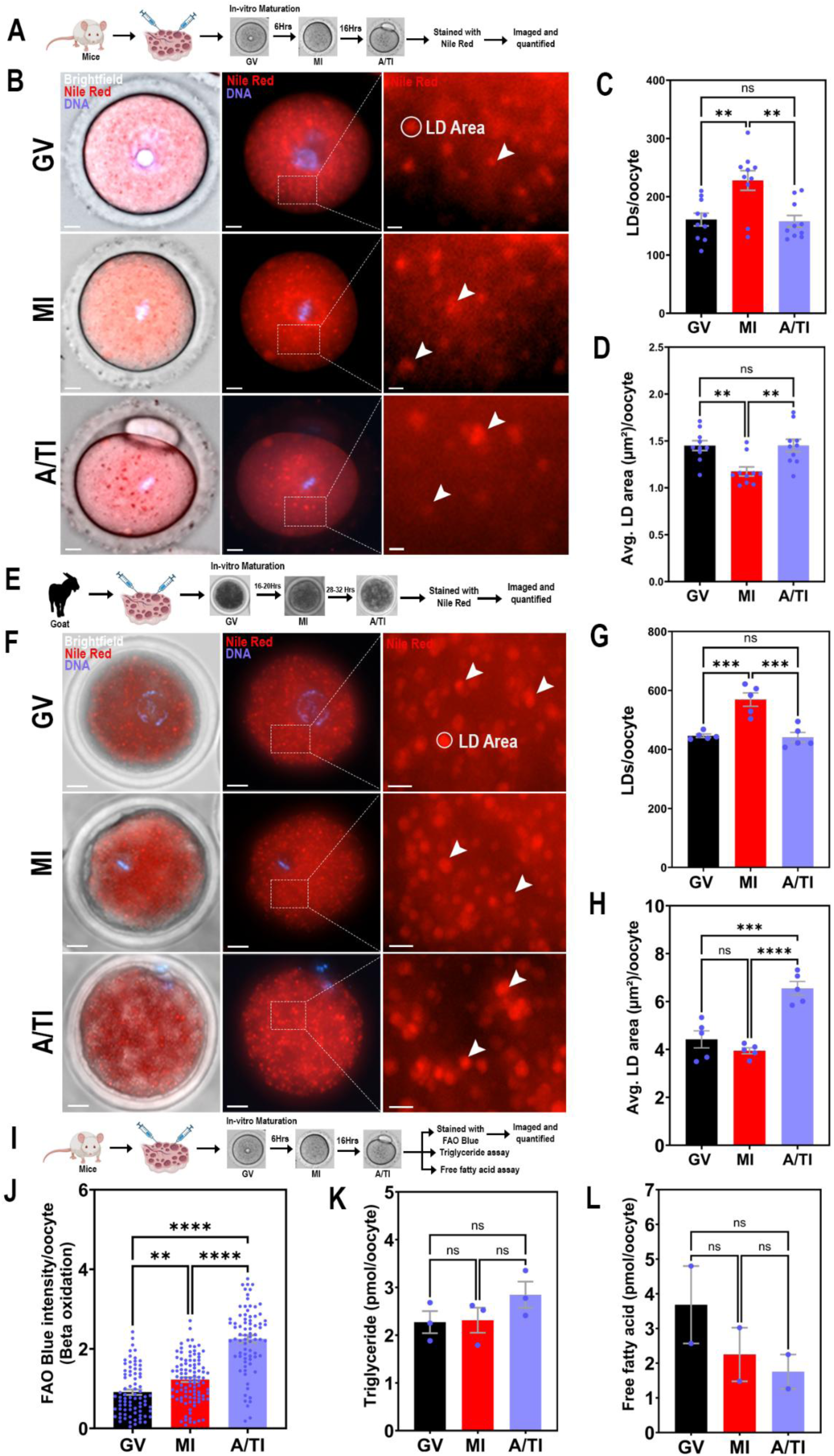
Dynamic remodeling of lipid droplets and metabolic pools during oocyte maturation in mouse and goat. **(A)** Schematic workflow for mouse oocyte in vitro maturation and Nile red staining. **(B)** Representative brightfield and fluorescence images of GV, MI, and A/TI oocytes stained with Nile Red (red) and Hoechst (DNA, blue). White arrowheads indicate LDs, and the white circle denotes the LD area. **(C–D)** Quantification of lipid droplet number per oocyte (C) and average lipid droplet area (D) in mouse oocytes. **(E)** Schematic workflow for goat oocyte in vitro maturation and Nile red staining. **(F)** Representative brightfield and fluorescence images of GV, MI, and A/TI goat oocytes stained with Nile Red (red) and Hoechst (blue). Arrowheads denote LDs, and a white circle marks the LD area used for quantification**. (G–H)** Quantification of lipid droplet number per oocyte (G) and average lipid droplet area (H) in goat oocytes. **(I)** Schematic workflow for oocyte lipid metabolic assays (FAO, triglycerides, free fatty acids). **(J)** Fatty acid oxidation (FAO) intensity per oocyte (20–30 oocytes per stage per replicate; total = 79 GV, 99 MI, and 77 MII oocytes across 3 independent biological replicates). **(K)** Quantification of Triglyceride and **(L)** Free fatty acid levels per oocyte in picomoles. For each experimental replicate, 100 oocytes were used for the TG and FFA assay, and values were normalized and expressed as per-oocyte levels in the graphs. Data are presented as mean ± SEM. Statistical significance was assessed using ordinary one-way ANOVA followed by Šídák’s multiple comparisons test. *p < 0.05, **p < 0.01, ***p < 0.001, ****p < 0.0001, ns = not significant. Scale bar: 10 µm; zoomed regions, 2 µm.

We next asked how LD dynamics relate to energy metabolism. Measurement of β-oxidation revealed a progressive rise from GV to MII (Fig. 1I, J), despite stable triglyceride (TG) and free fatty acid (FFA) levels (Fig. 1K, L). This paradox, elevated fatty acid catabolism without net depletion of lipid pools, suggests that oocytes maintain lipid homeostasis via active replenishment. Supporting this, we used C12 BODIPY, a fluorescent fatty acid analog that incorporates into newly formed LDs (Fig. 2A). GV-stage mouse oocytes were first stained with Nile Red to mark pre-existing LDs, then incubated with C12 BODIPY for 2 hours (Fig. 2B). Colocalization analysis revealed that all Nile Red-positive structures overlapped with C12 BODIPY signals, confirming that existing LDs persist during this period. Notably, additional C12 BODIPY-positive foci were observed that did not overlap with Nile Red, indicating active de novo LD biogenesis during maturation (Fig. 2B). This dynamic synthesis likely serves to replenish lipid reserves consumed during earlier stages of meiotic progression. To further identify the subcellular origins and associations of LDs, we performed colocalization studies using organelle-specific markers. MI-stage mouse oocytes were co-stained with ER-Tracker (endoplasmic reticulum), LysoTracker (lysosomes) and MitoTracker Green (mitochondria) (Fig. 2D-L). Quantitative analysis revealed that LDs predominantly colocalize with the endoplasmic reticulum, supporting its role as the primary site of LD biogenesis (Fig. 2E and F). LD association with lysosomes suggests that lipophagy may contribute to LD turnover, while mitochondrial interactions likely facilitate efficient lipid catabolism and energy production (Fig. 2G-L). Together, these data highlight the spatial and functional integration of LDs with key organelles to support energy homeostasis and redox balance during oocyte maturation.

**Fig 2:**
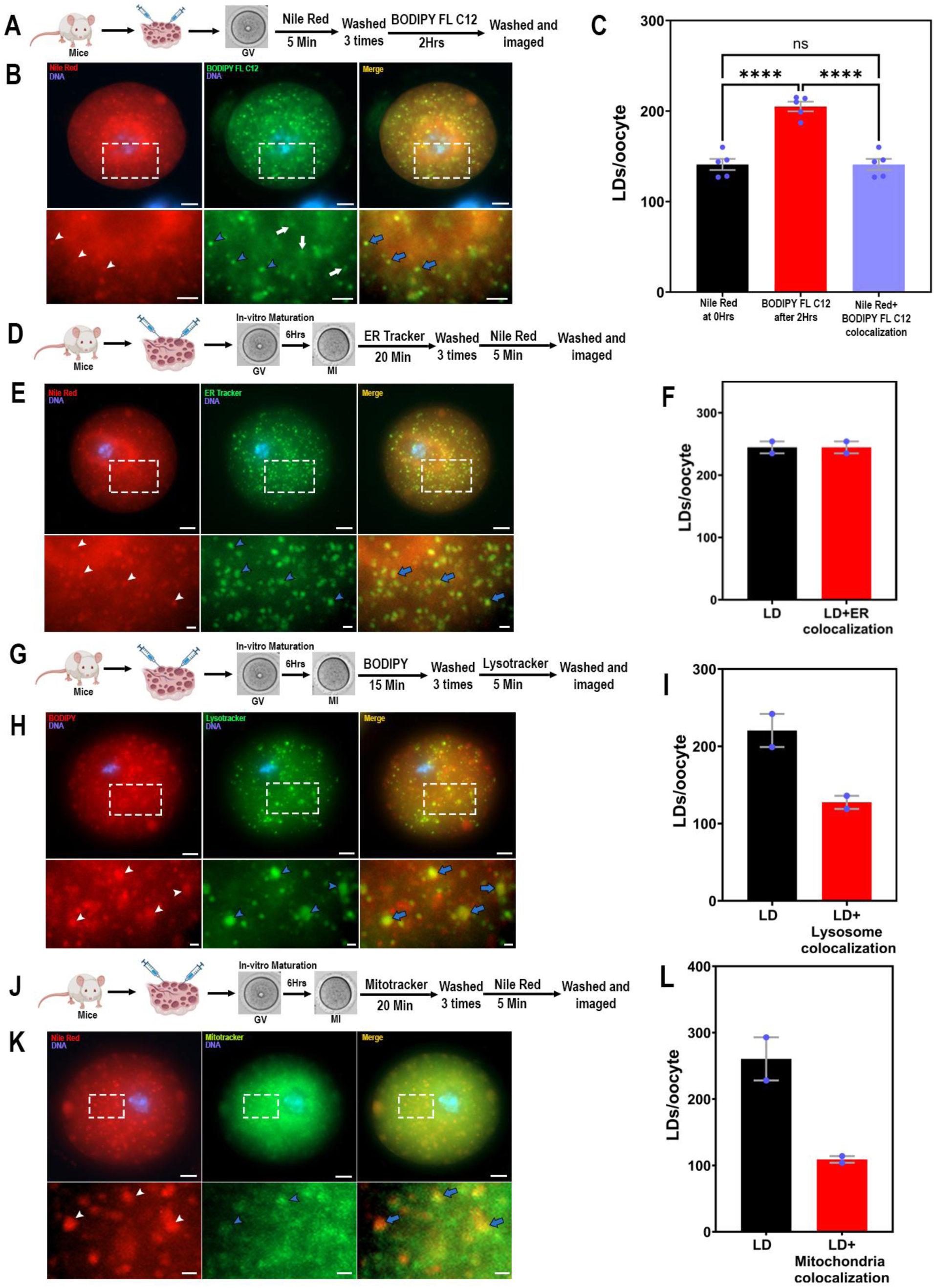
Oocyte lipid droplets incorporate de novo fatty acid synthesis and physically interact with ER, lysosomes, and mitochondria. **(A)** Schematic workflow for dual staining with Nile Red (LD marker) and BODIPY FL C12 (fatty acid tracer). **(B)** Representative fluorescence images of GV oocytes stained with Nile Red (red) marking pre-existing LDs and BODIPY FL C12 (green) marking de novo lipid synthesis. In zoomed regions (boxed areas), white arrowheads indicate pre-existing LDs, blue arrowheads indicate de novo synthesis within pre-existing LDs, white arrows denote newly synthesized LDs, and blue arrows highlight colocalization. **(C)** Quantification of lipid droplet number per oocyte after Nile Red, BODIPY FL C12, or colocalization. **(D)** Schematic workflow for Nile Red and ER Tracker labeling. **(E)** Representative images showing LDs (red) colocalized with ER (green). Zoomed regions highlight colocalization. **(F)** Quantification of LDs and LD–ER colocalization per oocyte. **(G)** Schematic Workflow for dual staining with BODIPY (LD marker) and LysoTracker (lysosome marker). **(H)** Representative images showing LDs (pseudo-red) colocalized with lysosomes (pseudo-green). Zoomed regions highlight colocalization. **(I)** Quantification of LDs and LD–lysosome Colocalization. (J) Schematic workflow for Nile Red and MitoTracker staining. **(K)** Representative images showing LDs (red) colocalized with mitochondria (green). Zoomed regions highlight colocalization. (L) Quantification of LDs and LD–mitochondria colocalization. In panels E, H, and K, white arrowheads indicate LDs, blue arrowheads indicate the respective organelles (ER, lysosomes, or mitochondria), and blue arrows indicate sites of colocalization. Data are presented as mean ± SEM. Statistical significance was assessed using ordinary one-way ANOVA followed by Šídák’s multiple comparisons test. (****) p < 0.0001; ns, not significant. Scale bar: 10 µm; zoomed regions in (B), 5 µm; other zoomed regions, 2 µm.

### DGAT1 is essential for lipid droplet biogenesis and meiotic progression

To dissect the functional role of LD biogenesis during oocyte maturation, we selectively inhibited diacylglycerol O-acyltransferase 1 (DGAT1) and DGAT2, enzymes catalyzing the final step in TG synthesis. DGAT1, predominantly localized to the ER, facilitates bulk TG production, while DGAT2, associated with LD surfaces, contributes to LD expansion and lipid storage (Fig. 3A and B). In mouse GV-stage oocytes, pharmacological inhibition of DGAT1 (40 µM) led to a marked reduction in LD formation, accompanied by decreased localization of DGAT1, perilipin-1 (PLIN1) and perilipin-2 (PLIN2) (Fig. 3C, D and Fig. S2. A). These two proteins are critical structural components of LDs that coat the droplet surface, regulate lipid metabolism, and protect stored lipids from lipolysis. Immunofluorescence analysis revealed fewer and less intense DGAT1, PLIN1, and PLIN2 puncta with DGAT1 inhibition, indicating compromised LD biogenesis and stability (Fig. S2A-G). Functionally, DGAT1 inhibition induced meiotic arrest at the MI stage in a significant proportion of oocytes (59.3 vs. 16.2% in control) as determined by DNA morphology, highlighting the essential role of DGAT1-mediated LD formation in supporting meiotic progression (Fig. 3E and Fig. S2H-J). Conversely, inhibition of DGAT2 at 20 µM had negligible effects on LD formation or meiotic maturation. Even at higher concentrations (150 µM), DGAT2 inhibition impaired maturation without affecting LD abundance, suggesting that DGAT2 may not be essential for de novo LD formation but may influence other aspects of lipid handling or oocyte physiology (Fig. S2K-N). These findings highlight distinct, non-redundant roles for DGAT1 and DGAT2 in oocyte lipid metabolism. To assess the evolutionary conservation of these mechanisms, we repeated the experiments in goat oocytes (Fig. 3J). Similar to the mouse, DGAT1 inhibition markedly reduced LD formation and meiotic maturation rates (6.7-fold vs. control) (Fig. 3K-M). Notably, however, DGAT1-inhibited goat oocytes displayed fewer but markedly larger LDs compared to controls, suggesting potential species-specific differences in LD regulation and turnover (Fig. 3K).

**Fig 3:**
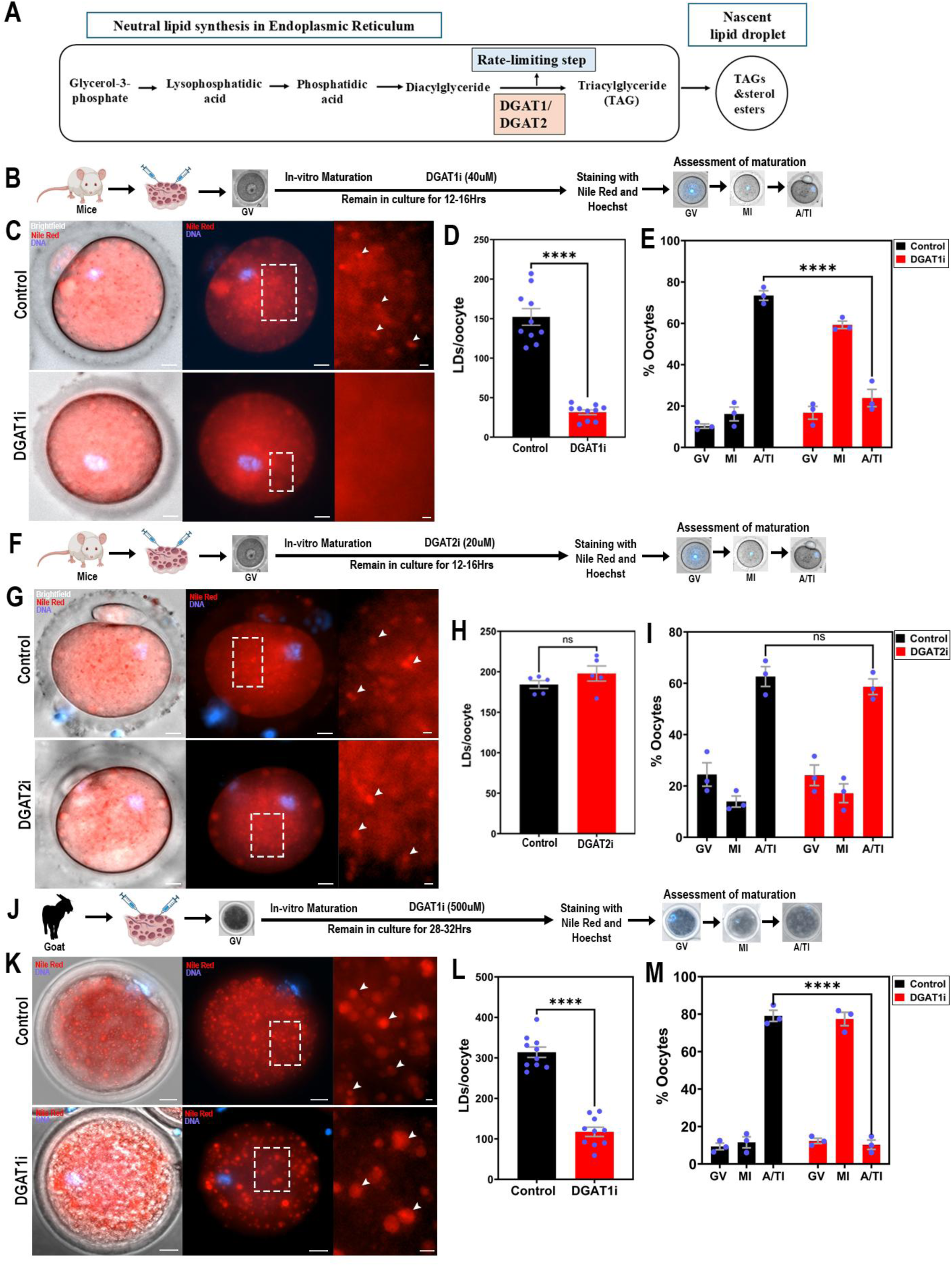
DGAT1 couples lipid droplet biogenesis to oocyte maturation across species. **(A)** Schematic of neutral lipid synthesis pathway in the endoplasmic reticulum leading to nascent lipid droplet formation. DGAT1/2 catalyze the rate-limiting step of triacylglyceride synthesis. **(B)** Schematic workflow showing DGAT1 inhibitor treatment during in vitro maturation (IVM), followed by staining and assessment of maturation. **(C)** Representative brightfield and fluorescence images of control and DGAT1i-treated oocytes stained with Nile Red (LDs, red) and Hoechst (DNA, blue). Zoomed regions highlight LDs (arrowheads). **(D)** Quantification of LD number per oocyte. **(E)** Percentages of oocytes at GV, MI, and A/TI stages (maturation assessment). **(F)** Schematic workflow showing DGAT2 inhibitor treatment during IVM, staining, and assessment of maturation. **(G)** Representative images of control and DGAT2i-treated oocytes. **(H)** Quantification of LD number per oocyte. **(I)** Percentages of oocytes at GV, MI, and A/TI stages (maturation assessment). **(J)** Schematic workflow showing DGAT1 inhibitor treatment in goat oocytes during IVM, followed by staining and maturation assessment. **(K)** Representative images of control and DGAT1i-treated goat oocytes stained with Nile Red (red) and Hoechst (blue). Zoomed regions highlight LDs (arrowheads). **(L)** Quantification of LD number per oocyte. **(M)** Percentages of oocytes at GV, MI, and A/TI stages (maturation assessment). (E, I, M) Oocyte maturation in mouse and goat oocytes (triplicate experiments, 25–35 oocytes per group per replicate). Data are presented as mean ± SEM. Statistical significance was assessed using unpaired t test (LD number, D, H, L) and two-way ANOVA with Šídák’s multiple comparisons (maturation, E, I, M). ****p < 0.0001; ns, not significant. Scale bars: mouse oocytes, 10 µm (zoomed 2 µm); goat oocytes, 20 µm (zoomed 5 µm).

### Loss of DGAT1 triggers metabolic overload, oxidative stress, and mitochondrial injury

Inhibition of DGAT1 impairs meiotic maturation, likely by disrupting lipid metabolic homeostasis and inducing lipotoxic stress in oocytes. DGAT1 inhibition significantly reduces LD formation, thereby limiting triglyceride storage and potentially increasing the availability of free fatty acids for β-oxidation. Consistent with this, quantification of free fatty acids and β-oxidation activity revealed a 3.1 and 2.9-fold increase in fatty acid availability and catabolism in DGAT1-inhibited oocytes, suggesting a compensatory metabolic shift to sustain energy production in the absence of sufficient lipid storage (Fig. 4A, B and C) (Fig. S3A, C). This elevated β-oxidation was accompanied by a significant rise in reactive oxygen species (ROS), as measured by DCFH-DA staining (1.25-fold increase compared to controls), indicating oxidative stress (Fig. 4D, E). Furthermore, mitochondrial membrane potential was notably reduced, suggesting compromised mitochondrial function (Fig. 4F, Fig. S3A, B). In parallel, cytoplasmic calcium levels were found to be increased, further implicating disrupted cellular homeostasis and stress signaling pathways (Fig. 4G, H).

**Fig 4:**
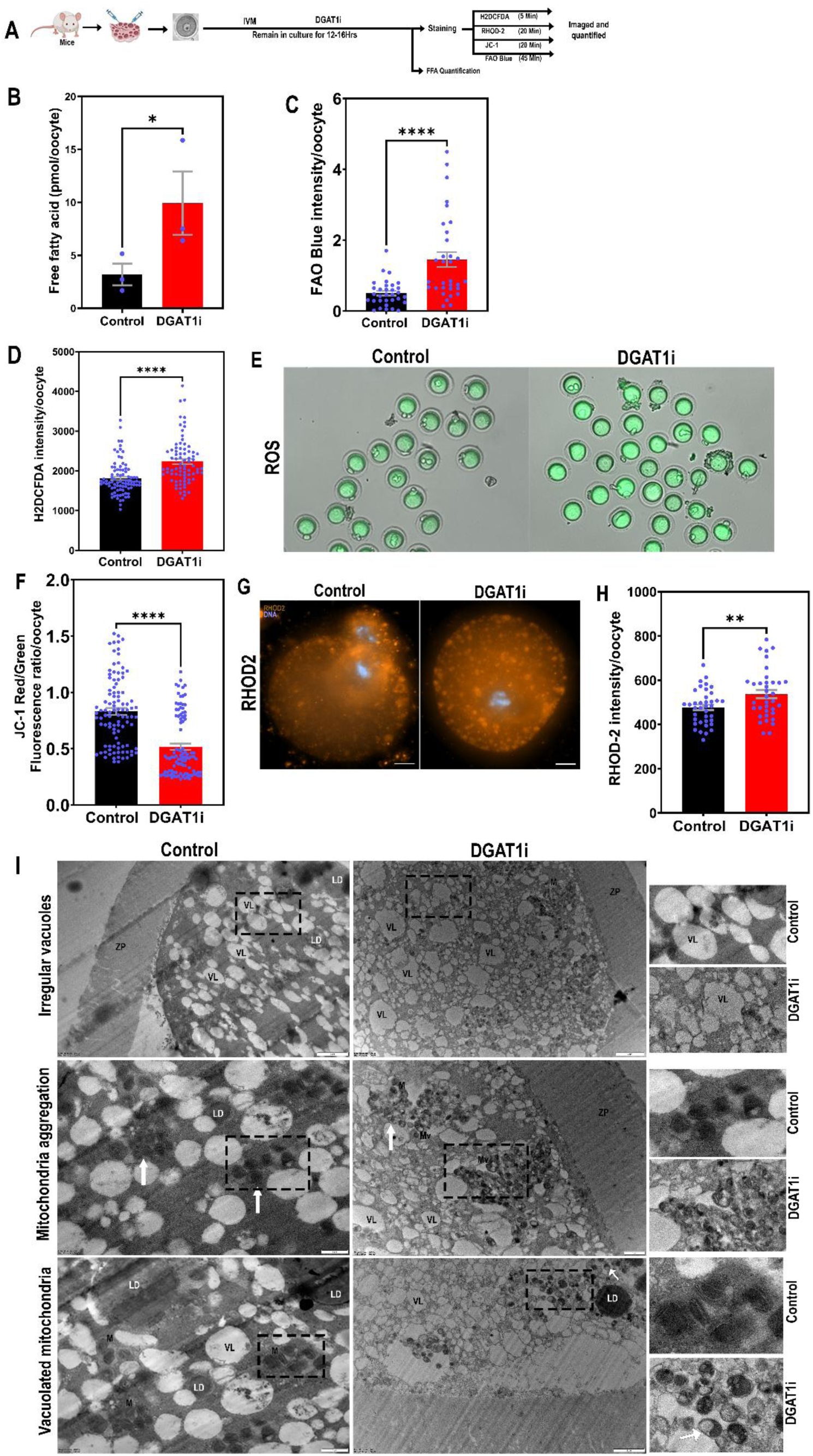
DGAT1 inhibition uncouples lipid storage from mitochondrial homeostasis, causing oxidative and structural stress in oocytes. **(A)** Schematic workflow for DGAT1 inhibitor treatment during in vitro maturation (IVM), followed by staining and FFA quantification. **(B)** Quantification of free fatty acids per oocyte. **(C)** Fatty acid oxidation (FAO) measured by FAO Blue fluorescence intensity. **(D)** Quantification of ROS by H2DCFDA staining. **(E)** Representative images of ROS fluorescence in control and DGAT1i-treated oocytes. **(F)** Quantification of mitochondrial membrane potential using JC-1 red/green fluorescence ratio. **(G)** Representative images of RHOD-2 fluorescence in control and DGAT1i-treated oocytes. **(H)** Quantification of mitochondrial calcium levels using RHOD-2 staining. **(I)** Transmission electron microscopy (TEM) images of control and DGAT1i-treated oocytes showing overall ultrastructure (normal view) and corresponding zoomed regions. DGAT1i-treated oocytes display ultrastructural abnormalities, including irregular vacuoles, mitochondrial aggregation, and vacuolated mitochondria. Abbreviations indicate: LD, lipid droplet; VL, vacuole; M, mitochondria; Mv, vacuolated mitochondria; ZP, zona pellucida. Data are presented as mean ± SEM. Statistical significance was assessed using one tailed unpaired t test: free fatty acids (*p < 0.05, N = 3 replicates with 100 oocytes each), FAO (****p < 0.0001, duplicate experiments; n = 32 control, 32 DGAT1i), ROS (****p < 0.0001, triplicate experiments; n = 88 control, 78 DGAT1i), Rhod-2 (**p < 0.01, duplicate experiments; n = 38 control, 36 DGAT1i), and JC-1 (****p < 0.0001, triplicate experiments; n = 100 control, 99 DGAT1i). Scale bars: fluorescence, 10 µm; TEM, 1–2 µm.

To gain ultrastructural insight into organelle integrity under DGAT1 inhibition, we performed transmission electron microscopy (TEM) on treated goat oocytes. TEM analysis revealed three major organelle abnormalities compared to controls: (1) irregular and disorganized vacuole structures, (2) aggregated mitochondria, and (3) mitochondria with extensive vacuolization (Fig. 4I). These structural defects are consistent with functional mitochondrial impairment and highlight the downstream consequences of lipid metabolic dysregulation. Together, these findings suggest that loss of DGAT1-mediated LD biogenesis may lead to metabolic overload, excessive β-oxidation, and ROS accumulation, culminating in mitochondrial damage and meiotic arrest. The observed organelle dysfunction and elevated oxidative stress underscore the essential role of de novo LD formation in maintaining metabolic and redox homeostasis.

### PANK2 is upregulated as a compensatory CoA biosynthetic response

To define the molecular programs underlying the metabolic stress induced by impaired LD biogenesis, we performed label-free quantitative proteomics on control and DGAT1-inhibited mouse oocytes. Across three independent replicates, proteome-wide profiling revealed a reproducible separation between the two conditions, as demonstrated by multivariate discriminant analysis and unsupervised hierarchical clustering (Fig. 5A–C). This separation indicates that inhibition of DGAT1 elicits a broad reprogramming of the oocyte proteome. Functional annotation of differentially expressed proteins revealed significant enrichment in categories linked to cytoplasmic structures (Fig. 5D). Biological process enrichment highlighted pronounced shifts in pathways governing lipid droplet organization, fatty acid catabolism, and oxidative stress responses (Fig. 5E). These categories aligned with our functional data, suggesting that loss of LD-derived triglyceride buffering forces oocytes to mobilize fatty acids excessively, fueling β-oxidation and escalating ROS burden. In parallel, molecular function analysis revealed significant enrichment in ATP hydrolysis, calcium channel activity, lipid-binding proteins, and enzymes involved in Coenzyme A (CoA) biosynthesis (Fig. 5F). This molecular signature pointed to elevated energy demand, altered ion flux, and activation of metabolic pathways supporting continuous fatty acid utilization. Among the top responders, Pantothenate Kinase 2 (PANK2), the rate-limiting enzyme in the CoA biosynthetic pathway, was markedly upregulated in DGAT1-inhibited oocytes (Fig. 5G). Immunolocalization confirmed this proteomic signature, revealing a 3.2-fold increase in PANK2 foci in the cytoplasm (Fig. 5H–J). Given its central role in generating CoA for fatty acid activation and mitochondrial β-oxidation, the induction of PANK2 likely represents a compensatory mechanism to maintain metabolic flux and redox buffering capacity when LD-dependent triglyceride storage is compromised. These findings position PANK2 as a critical metabolic node linking disrupted lipid droplet biogenesis to enhanced fatty acid catabolism and oxidative stress in mammalian oocytes.

**Fig 5:**
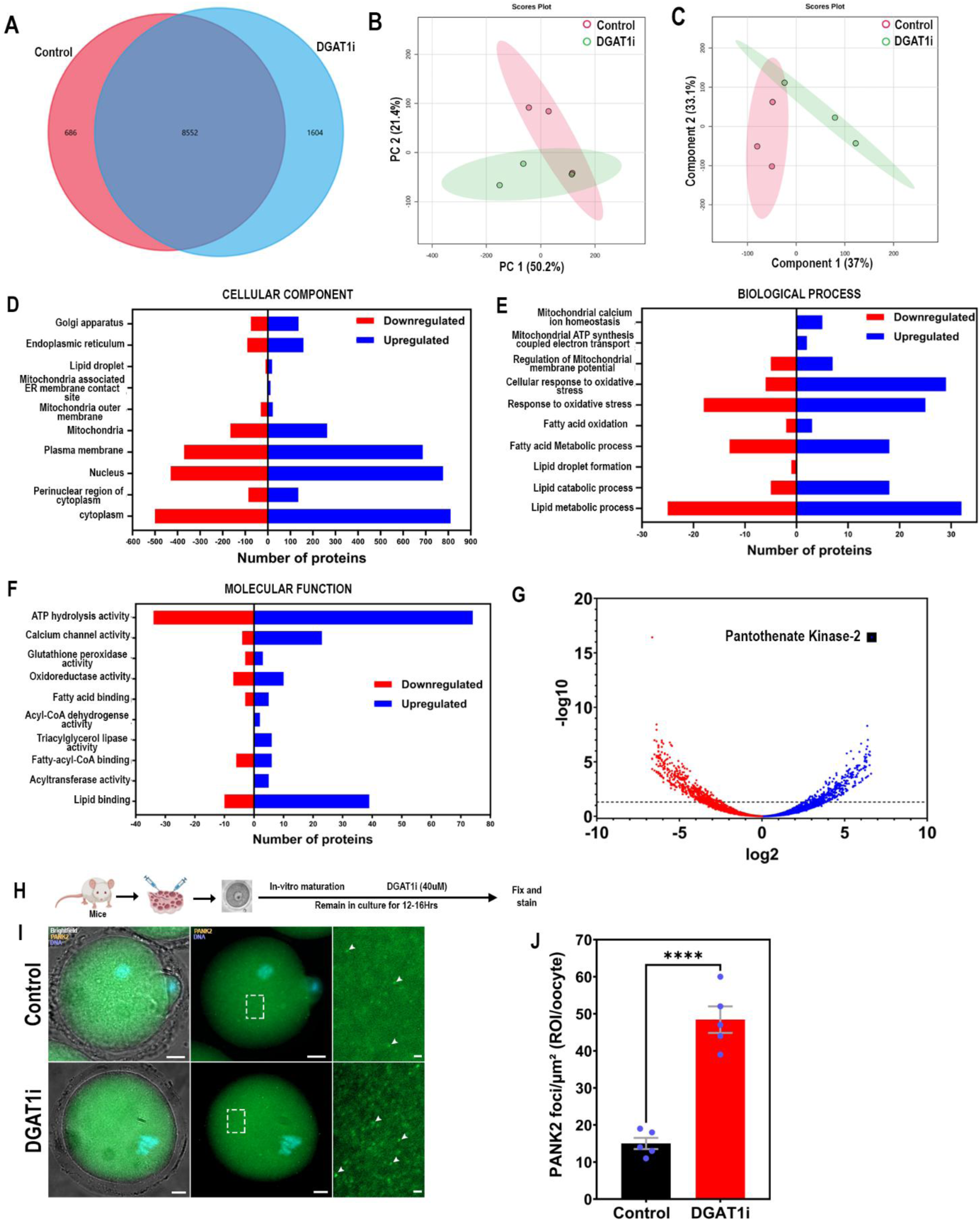
Proteomic profiling identifies PANK2 upregulation as a key metabolic response to DGAT1 inhibition in oocytes. **(A)** Venn diagram showing overlap of proteins identified in control and DGAT1i-treated oocytes proteome (n=3) **(B)** Principal component analysis (PCA) of proteomic profiles in control vs DGAT1i-treated oocytes. **(C)** Partial least squares discriminant analysis (PLS-DA) score plot of proteomic profiles in control vs DGAT1i-treated oocytes. **(D–F)** Gene ontology (GO) enrichment analysis of differentially expressed proteins in cellular components (D), biological processes (E), and molecular functions (F). Red = downregulated; blue = upregulated. **(G)** Volcano plot highlighting PANK2 as a significantly upregulated protein in DGAT1i-treated oocytes. **(H)** Schematic workflow of DGAT1i treatment during in vitro maturation followed by PANK2 immunostaining. **(I)** Representative fluorescence images of control and DGAT1i-treated oocytes stained for PANK2. White arrowheads indicate PANK2 foci, and zoomed regions highlight subcellular PANK2 foci. **(J)** Quantification of PANK2 foci per oocyte. Data are presented as mean ± SEM. Statistical significance was assessed using unpaired t test (****p < 0.0001). Staining was performed on 15 oocytes per group, of which 5 oocytes were quantified. Scale bars: fluorescence, 10 µm.

### PANK2 or β-oxidation inhibition alleviates oxidative stress and restores meiotic progression

Given the strong induction of PANK2 and the heightened β-oxidation signature upon DGAT1 inhibition, we hypothesized that excessive CoA-dependent fatty acid flux drives oxidative stress and meiotic arrest. To test this, we performed functional rescue experiments by pharmacologically inhibiting PANK2 or β-oxidation (Fig. 6A and B). Inhibition of PANK2 in DGAT1-inhibited oocytes resulted in a 2.38-fold increase in meiotic competence, with more oocytes progressing from GV through MI to MII compared with untreated controls. Concomitantly, ROS levels were 0.72-fold reduced, indicating that PANK2 suppression restores redox balance (Fig. 6C and Sup Fig.3D), suggesting that excessive PANK2 activity may contribute to oxidative stress and meiotic disruption when LD biogenesis is compromised. To further dissect the contribution of fatty acid catabolism, we inhibited mitochondrial β-oxidation using valproic acid (VPA). Similarly, VPA treatment rescued 2.03-fold meiotic competence (Fig. 6D). Thus, blocking β-oxidation prevents the oxidative burden imposed by excessive lipid utilization in DGAT1-deficient oocytes. Together, these rescue experiments establish that PANK2-driven CoA biosynthesis and β-oxidation act as dual effectors of oxidative stress and meiotic arrest under conditions of impaired LD biogenesis. Their inhibition partially restores meiotic progression and redox homeostasis, underscoring a critical interdependence between lipid droplet buffering, fatty acid oxidation, and oocyte quality under metabolic stress.

**Fig 6:**
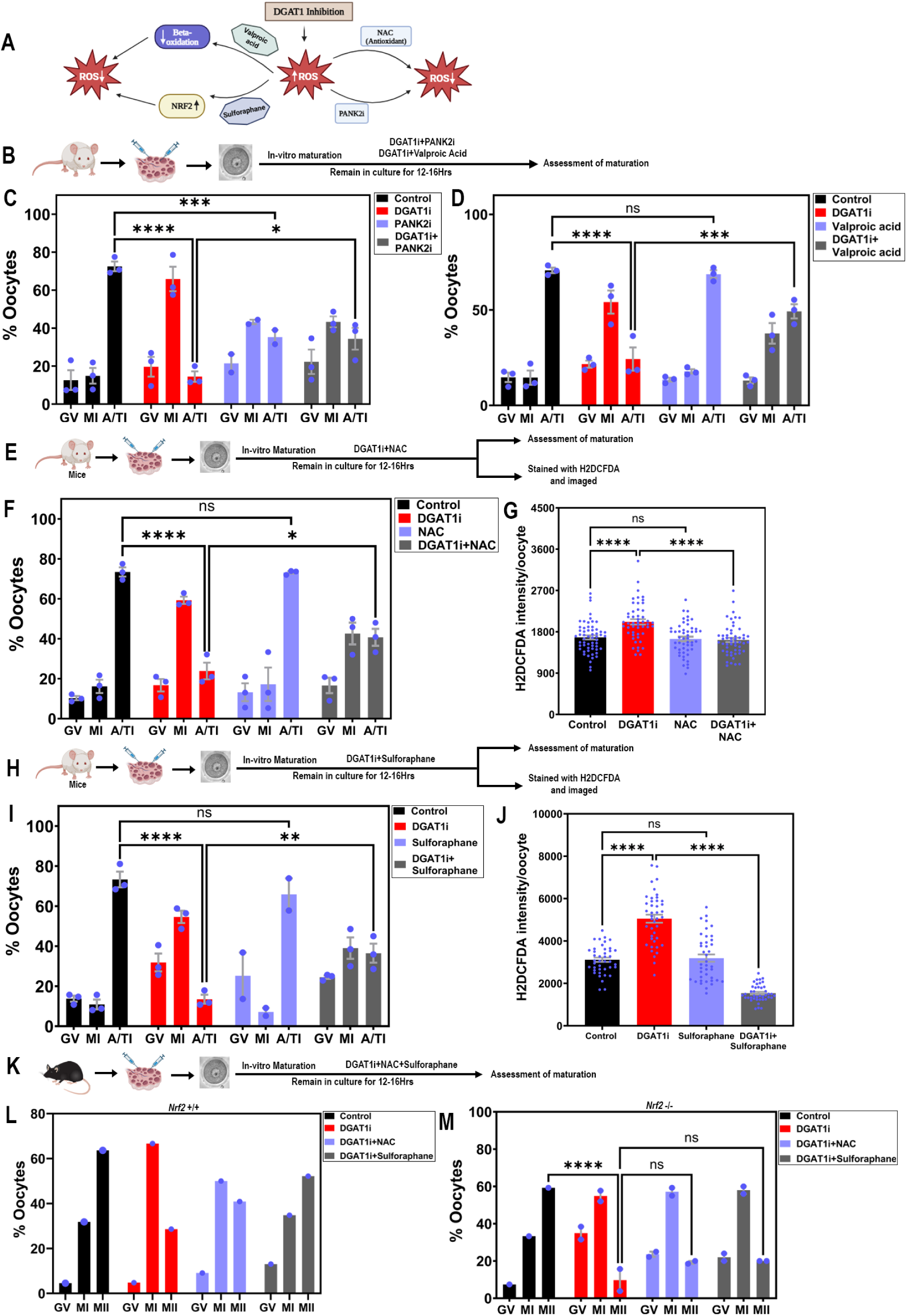
Antioxidant and NRF2-dependent interventions counteract DGAT1 inhibition–induced redox stress in oocytes. **(A)** Schematic model illustrating how DGAT1 inhibition elevates ROS and disrupts maturation, and how rescue agents (pantothenic acid, valproic acid, NAC, sulforaphane) or NRF2 signaling modulate this process. **(B)** Schematic workflow of treatments during in vitro maturation (IVM). **(C–D)** Percentage of oocyte maturation shown as percent oocytes at GV, MI, and A/TI stages after the indicated treatments. **(E)** Schematic workflow of DGAT1i±NAC treatment during IVM. **(F)** Percentage of oocyte maturation shown as percent oocytes at GV, MI, and A/TI stages. **(G)** ROS levels measured by H2DCFDA quantification per oocyte. **(H)** Schematic workflow of DGAT1i±Sulforaphane treatment during IVM. **(I)** Percentage of oocyte maturation shown as percent oocytes at GV, MI, and A/TI stages. **(J)** ROS levels measured by H2DCFDA quantification per oocyte. **(K)** Schematic workflow of treatments in *Nrf2*+/+ or *Nrf2*–/– oocytes during IVM. **(L–M)** Percentage of oocyte maturation shown as percent oocytes at GV, MI, and MII stages in wild-type (L) and *Nrf2*–/– (M) backgrounds. ROS assays were performed with 40–60 oocytes per condition. Maturation experiments were performed in triplicate with 25–35 oocytes per group per replicate. Data are presented as mean ± SEM. Statistical significance was assessed using ordinary one-way ANOVA or two-way ANOVA with Šídák’s multiple comparisons test. Statistical significance is indicated as follows: *p < 0.05; **p < 0.01; ***p < 0.001; ****p < 0.0001; ns = not significant.

### NRF2-dependent antioxidant signaling rescues meiotic competence

The results highlight the crucial role of redox balance, associated with fatty acid metabolism, in oocyte meiotic maturation. In addition, our proteomic analysis revealed that oxidative stress response pathways were altered in DGAT1-inhibited oocytes (Fig. 5), suggesting that defective redox homeostasis may contribute to meiotic arrest. To further investigate how oxidative stress influences meiotic competence and redox homeostasis in DGAT1-inhibited oocytes, a series of experiments involving the antioxidant N-acetylcysteine (NAC) and the NRF2 pathway were conducted (Fig. 6A-K). Nuclear factor erythroid 2-related factor 2 (Nrf2) is a crucial transcription factor that regulates cellular antioxidant defence mechanisms and plays a significant role in oxidative stress management. In response to oxidative stress, Nrf2 dissociates from its inhibitor, Keap1, translocates to the nucleus, and activates the expression of antioxidant and detoxifying enzymes such as superoxide dismutase (SOD) and glutathione peroxidase (GPx)^52^. In ovarian function, Nrf2 protects ovarian cells from oxidative damage, supports follicular development, and maintains oocyte quality^51^.

Initially, DGAT1-inhibited oocytes were treated with NAC to evaluate its ability to restore meiotic progression and redox balance (Fig. 6E). The results revealed that NAC treatment led to a 1.71-fold increase in meiotic progression, as evidenced by enhanced and progression to MII (Fig. 6F). Moreover, NAC treatment significantly reduced oxidative stress markers, including ROS levels, suggesting restoring redox balance in DGAT1-inhibited oocytes (Fig. 6G).

To confirm the involvement of NRF2 in this rescue, the same experiments were performed in NRF2 knockout (KO) oocytes. Remarkably, the NAC-induced restoration of meiotic progression was abolished in NRF2 KO oocytes, indicating that the NRF2 pathway is critical for the antioxidant effects of NAC. These results suggest that NRF2-mediated redox regulation is vital in restoring meiotic competence in DGAT1-inhibited oocytes. To further investigate the NRF2-redox balance, sulforaphane (SFN), a known NRF2 activator, was applied to DGAT1-inhibited oocytes (Fig. 6H). SFN treatment resulted in enhanced meiotic progression, improved GVBD and MII progression, and reduced oxidative stress markers (Fig. 6I, J). Similar to NAC, this enhancement in oocyte maturation was dependent on NRF2 activity, reinforcing the role of NRF2 in maintaining redox balance and meiotic competence under metabolic stress (Fig. 6 K, L). These experiments demonstrate that NRF2-dependent antioxidant treatment, through NAC and sulforaphane, can rescue meiotic progression and redox homeostasis in DGAT1-inhibited oocytes. The failure of this rescue in NRF2 KO oocytes underscores the essential role of NRF2 in regulating meiotic competence and mitigating oxidative stress.

### Aging disrupts de novo lipid droplet remodeling and DGAT1–ER coupling in oocytes

Because impaired LD biogenesis compromises meiotic progression in young oocytes, we next investigated whether age-associated decline in oocyte quality reflects similar defects in LD remodeling. Oocytes from reproductively young (2 months) and aged (14 months) females were compared (Fig. 7A). Quantitative imaging revealed that aged oocytes displayed a paradoxical phenotype: overall LD abundance was maintained or slightly elevated at the GV stage and showed little dynamic change during meiotic progression, yet mean LD area was significantly reduced (Fig. 7B–D). Instead of the large, remodeled droplets characteristic of young oocytes, aged oocytes contained fragmented, smaller LDs, pointing to defective remodeling and turnover.

**Fig 7:**
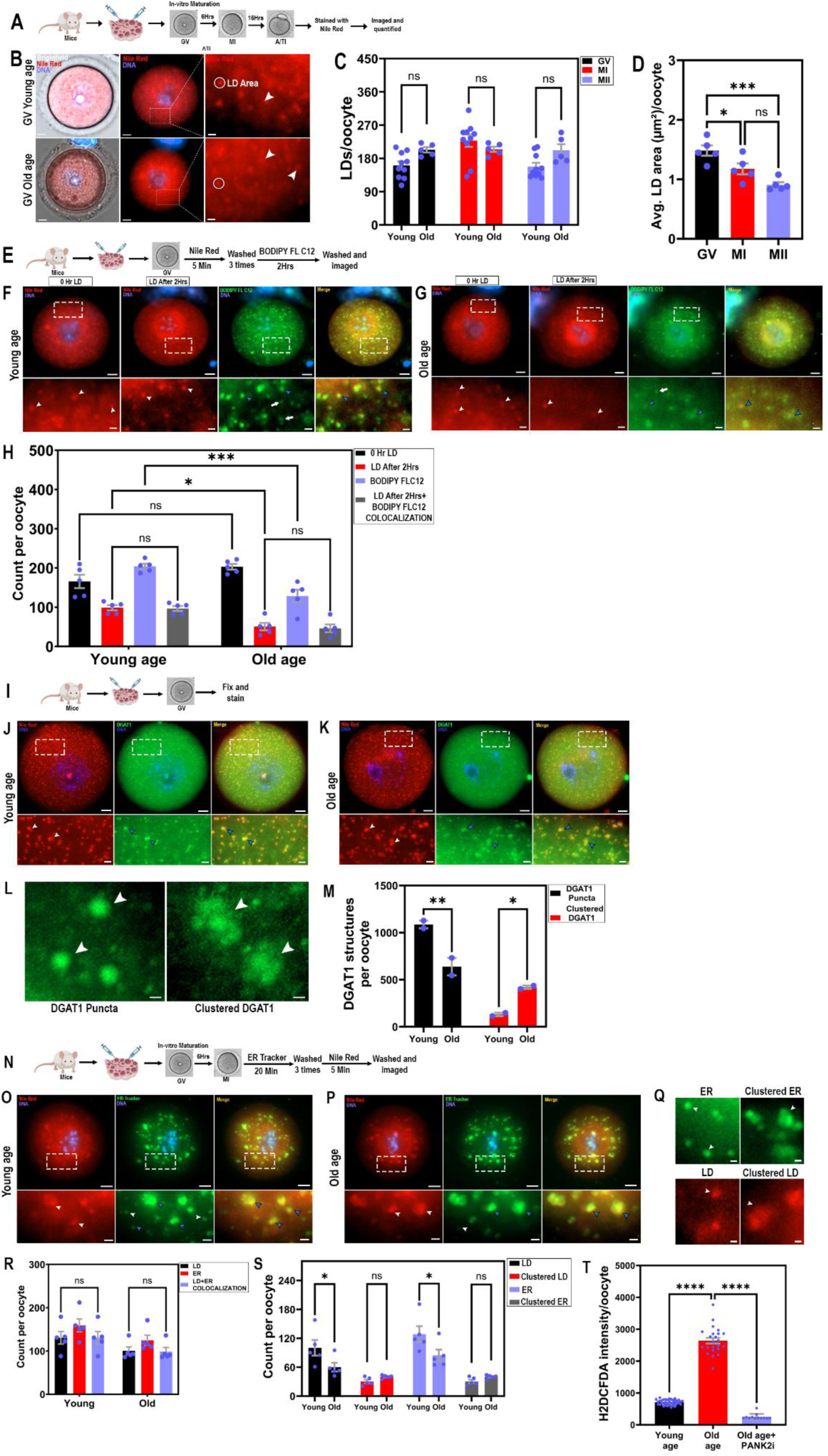
Age-driven disruption of lipid droplet biogenesis and metabolic cross-talk compromises oocyte redox homeostasis. **(A)** Schematic workflow for Nile Red staining of oocytes during in vitro maturation (IVM). **(B)** Representative brightfield and fluorescence images of GV-stage oocytes from young and old mice stained with Nile Red (red). The white circle indicates the LD area, and arrowheads show LDs. **(C)** Quantification of LD number per oocyte in young and old oocytes at GV, MI, and MII stages. **(D)** Average LD size per oocyte at GV, MI, and MII stages in old oocytes. **(E)** Schematic workflow for Nile Red and BODIPY FL C12 dual staining. **(F–G)** Representative images of GV-stage oocytes from young (F) and old (G) mice. Panel 1: Nile Red staining at 0 hr (pre-existing LDs); Panel 2: Nile Red staining after 2 hr; Panel 3: BODIPY FL C12 staining after 2 hr (de novo lipid synthesis); Panel 4: merged image showing colocalization. White arrowheads indicate pre-existing LDs, blue arrowheads indicate de novo synthesis within existing LDs and sites of colocalization, and white arrows indicate newly synthesized LDs. **(H)** Quantification of pre-existing LDs (Nile Red), newly synthesized LDs (BODIPY FL C12), and colocalized LDs**. (I)** Schematic workflow for DGAT1 immunostaining. **(J–K)** Representative images of GV-stage oocytes from young **(J)** and old **(K)** mice showing DGAT1 distribution (green) together with lipid droplets (LDs, red). Images were acquired at 60× magnification. Boxed areas are shown as zoomed regions. White arrowheads indicate LDs, and blue arrowheads indicate DGAT1 puncta/foci and their colocalization with LDs. **(L)** Additional zoomed images highlighting DGAT1 puncta and clustered DGAT1 foci. **(M)** Quantification of DGAT1 puncta and clustered DGAT1 per oocyte. **(N)** Schematic workflow for ER Tracker and Nile Red dual staining. **(O–P)** Representative images of GV-stage oocytes from young (O) and old (P) mice showing lipid droplet (LD, red) and endoplasmic reticulum (ER, green) interactions. Each panel contains three views: (1) LDs, (2) ER, and (3) merged colocalization. Boxed areas are shown as zoomed regions, where white arrowheads indicate LDs, white arrowheads in the ER panel indicate ER puncta, blue arrowheads indicate clustered ER, and blue arrowheads in the merged panel indicate LD–ER colocalization. **(Q)** Zoomed images highlighting ER puncta and clustered ER (upper panel) and LDs (lower panel). **(R)** Quantification of LD–ER colocalization in young and old oocytes. **(S)** Quantification of ER and LD puncta versus clustered structures in young and old oocytes. **(T)** Quantification of ROS intensity in young oocytes, old oocytes, and old oocytes treated with PANK2 inhibitor (PANK2i). Data are presented as mean ± SEM. Statistical significance was assessed using ordinary one-way ANOVA or two-way ANOVA with Šídák’s multiple comparisons test. Statistical significance is indicated as follows: *p < 0.05; **p < 0.01; ***p < 0.001; ****p < 0.0001; ns = not significant. Scale bars:10 µm (zoomed 2 µm); (L) 0.5 µm; (Q) 1 µm.

To test whether this phenotype reflected impaired de novo biogenesis, we performed metabolic labeling with C12-BODIPY. In young oocytes, tracer incorporation into LDs was robust throughout meiosis. In contrast, aged oocytes exhibited markedly reduced incorporation (Fig. 7E–H), indicating that although LDs accumulate with age, they are metabolically inert and not replenished through active biogenesis. This defect was consistent across mouse strains (Fig. S4A–G). (Fig. S4A-G). To mechanistically probe this defect, we examined DGAT1 localization. DGAT1 exhibited two spatial patterns in oocytes: discrete puncta and larger clusters. In aged oocytes, LDs continued to colocalize with DGAT1; however, there was a pronounced shift toward clustered DGAT1, suggesting misregulated enzyme organization at LD–ER interfaces (Fig. 7 I-M). A similar pattern was observed for ER–LD contacts: young oocytes exhibited balanced ER foci and clusters, whereas aged oocytes showed excessive ER clustering around LDs (Fig. 7N-S). This indicates that aging does not abolish DGAT1–LD or ER–LD contact formation, but instead skews their spatial architecture toward clustered assemblies that are likely dysfunctional.

Functionally, this structural imbalance correlated with elevated oxidative stress in aged oocytes, closely phenocopying the defects induced by pharmacological DGAT1 inhibition in young oocytes (Fig. 7T). Importantly, inhibition of PANK2 reduced ROS levels in aged oocytes, implicating maladaptive PANK2 activity in the oxidative imbalance (Fig. 7T). Together, these data establish that aging disrupts the DGAT1–ER axis of de novo LD biogenesis, resulting in excess but metabolically inert LD accumulation, defective remodeling, and increased oxidative stress. This mechanistic link between impaired LD remodeling and redox imbalance identifies defective de novo lipid biogenesis as a hallmark of reproductive aging.

## Discussion

### Energy homeostasis mediated by lipid droplets

Dynamic changes in LDs abundance and size during meiotic progression underscore their critical role in meeting the energy demands of oocyte maturation. The observed increase in LD abundance during the GV to MI transition, followed by a decline at MII, reflects a finely tuned process of lipid mobilization. This mobilization supports energy-intensive processes such as spindle assembly, chromosome segregation, and cytoskeletal reorganization. Elevated levels of β-oxidation observed during the GV to MII stages further confirm the coordinated use of lipid reserves to efficiently generate ATP during these critical meiotic transitions. De novo LD synthesis, visualized through C12 BODIPY staining, highlights an adaptive response by oocytes to replenish lipid reserves depleted during meiotic progression.

LDs are indispensable for oocyte function despite the availability of carbohydrates and lipids as direct energy sources. As metabolically active cells, oocytes require substantial energy to support meiotic progression, spindle assembly, and early embryonic development. LDs act as intracellular reservoirs of neutral lipids, predominantly triglycerides, which are hydrolyzed into free fatty acids (FFAs) for mitochondrial β-oxidation. This pathway is highly efficient in producing ATP, ensuring a steady energy supply during the energetically demanding phases of oocyte maturation.

In addition to energy production, LD dynamics play a pivotal role in redox homeostasis. Fatty acid metabolism via β-oxidation generates ROS. LDs mitigate oxidative stress by sequestering oxidized lipids, thus shielding the oocyte from ROS-induced damage. This dual role in energy generation and ROS buffering underscores the importance of LDs in maintaining oocyte quality and developmental competence. The association of LDs with the ER and lysosomes, observed in this study, emphasizes their multifaceted role in lipid turnover and oxidative stress management through processes like lipophagy. This interplay equips oocytes to handle fluctuating metabolic and oxidative demands during maturation. Disruptions in LD biogenesis or turnover, such as excessive lipolysis or abnormal lipid accumulation, can upset this balance, resulting in impaired meiotic progression, heightened oxidative damage, and diminished developmental competence.

### DGAT1 as a central regulator of lipid droplet biogenesis and oocyte competence

Our findings establish DGAT1 as a key regulator of de novo LD biogenesis, which is essential for coordinating metabolic and redox homeostasis during oocyte maturation. Pharmacological inhibition of DGAT1 led to profound defects in LD formation, resulting in meiotic arrest at the GV or MI stages. This arrest was accompanied by a metabolic crisis characterized by elevated β-oxidation, increased ROS, reduced mitochondrial membrane potential, and dysregulated intracellular calcium. Ultrastructural analysis revealed organelle damage, including vacuolated and aggregated mitochondria and disrupted vacuole morphology, highlighting the downstream consequences of impaired lipid storage and buffering capacity. Intriguingly, germline DGAT1 knockout mice remain fertile, albeit with reduced litter sizes and overall fecundity. This observation raises an important mechanistic question: why does acute pharmacological inhibition of DGAT1 in oocytes result in meiotic failure, whereas complete genetic ablation does not completely abolish fertility?

One plausible explanation is metabolic compensation in the germline DGAT1 knockout model, where chronic DGAT1 loss throughout development allows time for adaptive responses, such as upregulation of DGAT2 or alternative lipid metabolic pathways, to partially restore LD homeostasis in oocytes. Indeed, developmental plasticity may enable oocytes in knockout mice to maintain a minimal threshold of lipid storage and energy buffering capacity sufficient to support maturation in a subset of cells, albeit with diminished efficiency. By contrast, acute pharmacological inhibition of DGAT1 in isolated GV-stage oocytes likely induces a sudden metabolic blockade, preventing the activation of compensatory pathways. The oocyte’s limited transcriptional and translational capacity at this stage may further constrain its ability to mount an effective adaptive response, leading to unbuffered lipid dysregulation, ROS accumulation, and meiotic failure. This distinction between chronic adaptation versus acute inhibition underscores the vulnerability of the oocyte’s metabolic machinery and suggests that the timing and context of DGAT1 function are critical determinants of meiotic competence. Interestingly, DGAT2 inhibition had minimal impact on oocyte maturation or LD formation, reinforcing the specificity of DGAT1 as the principal enzyme mediating triglyceride synthesis and LD biogenesis during meiotic progression. This functional divergence highlights non-redundant roles of DGAT1 and DGAT2 in oocyte lipid metabolism.

### PANK2 and NRF2 define adaptive but insufficient stress responses

Proteomic analysis uncovered PANK2 upregulation as a key compensatory response to DGAT1 inhibition, consistent with an attempt to sustain CoA biosynthesis and drive fatty acid oxidation. However, this adaptation exacerbated oxidative stress, linking excessive β-oxidation to meiotic failure. Pharmacological inhibition of PANK2 or β-oxidation partially restored meiotic competence and redox balance, highlighting their maladaptive role in the absence of adequate LD buffering. Concurrently, NRF2 signaling emerged as the central antioxidant safeguard. Both N-acetylcysteine supplementation and sulforaphane-mediated NRF2 activation restored meiotic progression in DGAT1-deficient oocytes, but these benefits were abolished in NRF2 knockout oocytes. Together, these data establish a DGAT1–PANK2–NRF2 axis that coordinates lipid storage, fatty acid utilization, and redox homeostasis to preserve oocyte quality.

### Aging as a failure of lipid droplet remodeling

Metabolic labeling with C12-BODIPY revealed that while young oocytes robustly incorporated tracer into LDs during meiotic maturation, aged oocytes showed markedly diminished incorporation. Thus, despite an increase in LD number with age, these structures become metabolically inert and are not replenished through active biogenesis. This defect was strain-independent and associated with altered DGAT1 organization. Whereas DGAT1 normally localizes as both discrete puncta and smaller clusters at LD–ER contact sites, aged oocytes exhibited a pronounced shift toward clustered DGAT1 and ER, suggesting that aberrant enzyme distribution underlies defective LD remodeling. A recent study using label-free chemical imaging of human oocytes similarly reported age-dependent changes in the lipid fingerprint, including increases in LD size and number, accumulation of oxidized lipids and carotenoids, and structural evidence of membrane and mitochondrial degradation ^57^. These human findings parallel our observations: LDs accumulate but lose metabolic plasticity, while oxidative stress signatures emerge. Our work provides a mechanistic framework in which the DGAT1–PANK2–NRF2 axis coordinates de novo LD remodeling, fatty acid flux, and redox buffering to preserve oocyte fitness. Disruption of this axis, whether through aging or enzyme mislocalization, results in failed LD remodeling, maladaptive lipid oxidation, mitochondrial stress, and impaired meiotic progression. Together, these results suggest that the DGAT1–PANK2–NRF2 pathway acts as a conserved safeguard of oocyte quality, and its failure is a central feature of reproductive aging across species.

### Therapeutic implications and future directions

The identification of DGAT1 as a key regulator of lipid metabolism in oocytes offers promising therapeutic opportunities to enhance fertility outcomes. Strategies to augment LD biogenesis or modulate DGAT1 activity may counteract oxidative stress and energy imbalances in aging or metabolically compromised oocytes. Additionally, targeting NRF2 signaling pathways could further bolster redox homeostasis, complementing interventions aimed at preserving oocyte integrity. Future investigations should focus on the molecular interactions between DGAT1-mediated LD formation, mitochondrial function, and redox signaling pathways such as NRF2. Moreover, exploring the roles of other lipid metabolism enzymes, including DGAT2, under varying metabolic conditions, may provide a comprehensive understanding of their contributions to oocyte maturation. Translating these findings to human oocytes could have significant implications for improving assisted reproductive technologies (ART).

In summary, we establish lipid droplets as dynamic metabolic organelles that couple fatty acid utilization with redox buffering to ensure meiotic fidelity. DGAT1-mediated de novo LD biogenesis is essential for this process, and its disruption, either experimentally or with age, triggers maladaptive PANK2-driven β-oxidation, oxidative stress, and meiotic failure. These findings reveal defective LD remodeling as a hallmark of reproductive aging and highlight the DGAT1–PANK2–NRF2 axis as a mechanistic foundation for future therapeutic strategies in fertility preservation.

## Materials and methods

### Animal ethics

FVB mice (7–9 weeks old) were primarily used for all experiments. For age-comparison studies, both young (7–9 weeks old) and aged (16 months old) FVB mice, as well as young (7–9 weeks old) and aged (13 months old) C57BL/6J mice, were included. In addition, B6.129X1-Nfe2l2^tm1Ywk/J (Nrf2 knockout) mice (2 months old) and Nrf2 wild-type mice of the same background were used for selected experiments. All animals were maintained under a 12 h light/12 h dark cycle at a constant temperature of 21–22 °C, with food and water provided ad libitum, in accordance with the guidelines of the Animal Ethics Committee of the National Institute of Animal Biotechnology (NIAB).

### Mice oocyte Collection and In-Vitro Maturation

Mice were euthanized, and ovaries were surgically excised and transferred into 1 mL of pre-warmed M2 medium. Oocytes were released by carefully puncturing the ovaries with a fine needle and collected into M2 medium. They were washed four to five times in 100 µL M2 medium drops to remove debris. For in vitro maturation, groups of at least 20–30 oocytes were cultured in 100 µL drops of maturation medium covered with mineral oil in a 35 mm culture dish and incubated at 37 °C in a CO₂ incubator under the specified treatment or control conditions. For experiments requiring specific stages, oocytes were collected at defined time points: germinal vesicle (GV) stage at 0 h, metaphase I (MI) stage after 6 h, and metaphase II (MII) stage after 12–16 h of culture.

### Goat Oocyte Collection and In-Vitro Maturation (IVM)

Ovaries from Osmanabadi goats were obtained from a local slaughterhouse and transported to the laboratory in 1× PBS. Cumulus–oocyte complexes (COCs) were aspirated from antral follicles using an 18-gauge needle in TCM-199 medium. Only oocytes surrounded by 3–4 layers of compact cumulus cells were selected for further experiments.

Selected oocytes were washed thoroughly to remove debris and cultured in 100 µL drops of IVM medium containing TCM-199, 4 IU/mL follicle-stimulating hormone (FSH), 4 IU/mL luteinizing hormone (LH), 1 µg/mL Estradiol, 10% follicular fluid, 10% fetal bovine serum (FBS), 0.81 mM sodium pyruvate, 10 ng/mL EGF, and 50 µg/mL gentamycin. Cultures were maintained under mineral oil at 38.5 °C in a humidified CO₂ incubator (5% CO₂) for 28–32 h.

For stage-specific experiments, oocytes were collected at defined time points: germinal vesicle (GV) stage (0 h), metaphase I (MI) stage (∼16 h), and metaphase II (MII) stage (28–32 h).

### Staining

#### Lipid Droplet Staining with Nile Red

A stock solution of Nile Red (SRL, Cat. No. 47353) was prepared by dissolving 1 mg of dye in 1 mL of DMSO to obtain a 1 mg/mL stock solution. A 10 µg/mL working solution was freshly prepared by diluting the stock in culture medium. Oocytes were incubated in the working solution for 5 min at 37 °C in a CO₂ incubator. After incubation, oocytes were washed three times with fresh medium to remove excess dye. Imaging was performed using the AF568 channel on a Zeiss Axio Observer.Z1/7 microscope.

#### Lipid Droplet Staining with BODIPY

A stock solution of BODIPY [((3,5-dimethyl-1H-pyrrol-2-yl)(3,5-dimethyl-2H-pyrrol-2-ylidene)methyl)methane] (difluoroborane) (TCI, Cat. No. D5724) was prepared by dissolving 1 mg of dye in 1 mL of DMSO to obtain a 1 mg/mL stock solution. A 10 µg/mL working solution was freshly prepared by diluting the stock in culture medium. Oocytes were incubated in the working solution for 20 min at 37 °C in a CO₂ incubator. After incubation, oocytes were washed three times with fresh medium to remove excess dye. Fluorescence imaging was performed using the AF488 channel on a Zeiss Axio Observer.Z1/7 microscope.

#### Fatty Acid Tracing with BODIPY™ FL C12

Fatty acid uptake and de novo lipid synthesis were assessed using BODIPY™ FL C12 (ThermoFisher, Cat. No. D3822). A 5 mM stock solution was prepared in DMSO and stored at –20 °C in aliquots. An intermediate stock of 0.5 mM was prepared in DMSO by 1:10 dilution of the main stock. For experiments, a 5 µM working solution was freshly prepared by diluting the intermediate stock 1:100 in culture medium. Oocytes were incubated in the working solution for 2 h at 37 °C in a CO₂ incubator. After incubation, oocytes were washed three times with fresh medium to remove unincorporated dye. Fluorescence imaging was performed using the AF488 channel on a Zeiss Axio Observer.Z1/7 microscope.

#### Endoplasmic Reticulum Staining with ER-Tracker™ Green

A 1 mM stock solution of ER-Tracker™ Green (Cat. No. E34251, ThermoFisher) was prepared in DMSO. Oocytes were incubated in culture medium containing 5 µM ER-Tracker™ Green for 20 min at 37 °C in a CO₂ incubator, followed by three washes with fresh medium. Imaging was performed using the AF488 channel.

#### Lysosomal staining with Lysotracker™ Red

**A** 10 µM stock solution of Lysotracker™ Red (Cat. No. L7528, ThermoFisher) was prepared in PBS. For staining, 2 µL of stock solution was added to 100 µL of culture medium to achieve a final concentration of 0.2 µM. Oocytes were incubated for 5 min at 37 °C in a CO₂ incubator, followed by three washes with fresh medium. Imaging was performed using the AF555 channel.

#### Mitochondrial staining with MitoTracker™ Green FM

A 5 µM stock solution of MitoTracker™ Green FM (Invitrogen, Cat. No. M7514) was prepared in DMSO. For staining, a 200nM working solution was freshly prepared by diluting the stock in culture medium. Oocytes were incubated in the working solution for 20 min at 37 °C in a CO₂ incubator. After incubation, oocytes were washed three times with fresh medium to remove unbound dye. Fluorescence imaging was performed using the AF488 channel on a Zeiss Axio Observer.Z1/7 microscope.

#### Mitochondrial membrane potential staining with JC-1

A stock solution of JC-1 (Invitrogen, Cat. No. M34152) was prepared in DMSO (100 µM). For staining, oocytes were incubated with 2 µM final JC-1 for 30 min at 37 °C in a CO₂ incubator. After incubation, oocytes were washed three times with fresh medium before imaging. Fluorescence was acquired on a Zeiss Axio Observer.Z1/7 microscope using 488 nm excitation for JC-1 monomers (green channel) and 555 nm excitation for JC-1 aggregates (red channel).

#### Fatty acid β-oxidation detection with FAO Blue

A stock solution of FAO Blue (Fatty Acid Oxidation Detection Reagent; Diagnocine, Cat. No. FNK-FDV-0033) was prepared at 0.5mM in DMSO. For staining, a 20 µM working solution was freshly prepared by diluting the stock in DPBS. Since intact oocytes exhibited poor dye permeability, a brief Pronase treatment was performed prior to staining. Specifically, 2 µL of a 10 mg/mL Pronase stock was added to 100 µL of culture medium containing oocytes and incubated for 2 min at 37 °C, followed by immediate washing with fresh medium to stop enzymatic activity.

After permeabilization, oocytes were incubated with the 20 µM FAO Blue working solution in DPBS for 45 min at 37 °C in a CO₂ incubator. To account for the intrinsic auto fluorescence of oocytes at 405 nm excitation, images were acquired both before and after FAO Blue staining, and dye-specific signals were subsequently analysed.

#### Mitochondrial Calcium Staining with Rhod-2 AM

Mitochondrial calcium was detected using Rhod-2 AM, cell-permeant (Invitrogen, Cat. No. R1245MP). A 500 µM stock solution was prepared in DMSO and diluted in culture medium to obtain a 5 µM working solution. Oocytes were incubated in 100 µL of the working solution for 40 min at 37 °C in a CO₂ incubator. After incubation, oocytes were washed three times with fresh medium to remove excess dye. Fluorescence imaging was performed using the AF555 channel on a Zeiss Axio Observer.Z1/7 microscope.

#### ROS Detection with H₂DCFDA

Reactive oxygen species (ROS) levels in oocytes were assessed using the fluorescent probe H₂DCFDA (2′,7′-dichlorodihydrofluorescein diacetate; Invitrogen, Cat. No. D399). Oocytes were incubated in 10 µM H₂DCFDA diluted in maturation medium for 5 min at 37 °C.

Following incubation, oocytes were washed with fresh medium to remove excess dye and immediately imaged under a fluorescence microscope using the Alexa Fluor 488 channel.

### Treatments

#### Lipid Droplet Inhibition During Oocyte Maturation

To inhibit lipid droplet formation, oocytes were treated with DGAT1 and DGAT2 inhibitors during in vitro maturation.

For DGAT1 inhibition, T863 (DGAT1 inhibitor; Sigma-Aldrich, Cat. No. SML0539) was used. A 4 mM stock solution was prepared in DMSO and stored at –20 °C. For mouse oocytes, a 40 µM working solution was freshly prepared in maturation medium, and oocytes were incubated in 100 µL of inhibitor-containing medium for 12–16 h at 37 °C in a CO₂ incubator. A lower concentration (20 µM) was initially tested and showed only a slight reduction in maturation efficiency and slight reduction in lipid droplets. therefore, 40 µM was used for all subsequent mouse experiments. For goat oocytes, higher concentrations were tested; a phenotype was observed at 500 µM, and additional trials were performed at 100 µM and 200 µM.

For DGAT2 inhibition, PF-06424439 (DGAT2 inhibitor; Sigma-Aldrich, Cat. No. PF-06424439) was used. A stock solution was prepared in DMSO and stored at –20 °C. Working solutions were freshly prepared in maturation medium at final concentrations of 20 µM, 50 µM, 100 µM, and 150 µM. Oocytes were incubated in 100 µL of inhibitor-containing medium for 12–16 h at 37 °C in a CO₂ incubator.

In both cases, vehicle control oocytes were treated with the corresponding concentration of DMSO.

#### Rescue of DGAT1 Inhibition–Induced Arrest

To assess whether DGAT1 inhibitor–induced oocyte maturation arrest could be rescued, pharmacological agents were co-administered with T863 during in vitro maturation. Oocytes were cultured in 100 µL of maturation medium containing 40 µM T863 together with the respective rescue agent for 12–16 h at 37 °C in a CO₂ incubator. Vehicle controls (DMSO or sterile water, matched to the solvent for each compound) were included in all experiments. Additional controls included oocytes treated with inhibitor alone, rescue agent alone, or vehicle.

#### N-Acetyl-L-cysteine (NAC)

NAC (Sigma-Aldrich, Cat. No. A9165) was tested at 100, 250, and 500 µM. The 250 µM concentration partially rescued the DGAT1 inhibitor–induced phenotype.

#### Valproic Acid

Valproic acid (MedChemExpress, Cat. No. HY-10585) was tested at 250, 300, and 750 µM. The 250 µM concentration partially rescued the phenotype.

#### Pantothenate Kinase Inhibitor (PANK2i)

Pantothenate Kinase-IN-2 (MedChemExpress, Cat. No. HY-44170) was tested at 10, 20, 40, and 100 µM. The 40 µM concentration partially rescued the phenotype.

#### Sulforaphane

Sulforaphane (MedChemExpress, Cat. No. HY-13755) was tested at 0.25 µM. This concentration partially rescued the DGAT1 inhibitor–induced phenotype.

#### Immunocytochemistry

Oocytes were fixed in 4% paraformaldehyde (PFA) for 30 min in a 100 µL drop and washed three times with 0.5% TBST (TBS containing 0.5% Tween-20). Permeabilization was performed with 0.1% Triton X-100 for 5 min in a 100 µL drop, followed by washing three times with 0.5% TBST. Non-specific binding was blocked using 10% ADB (antibody dilution buffer) for 15 min (twice) in 100 µL medium. After blocking, oocytes were incubated overnight at 4 °C in a humidified chamber with primary antibodies against PLIN1 (Cat# A4758, Abclonal), PLIN2 (Cat# A22456, Abclonal), DGAT1 (Cat# A6857, Abclonal), and PANK2 (Cat# 11001-1-AP, Proteintech), each diluted 1:100 in full-strength ADB. Following incubation, oocytes were washed three times with 0.5% TBST (5 min each), then blocked again with 10% ADB for 15 min (twice). Oocytes were subsequently incubated with the appropriate fluorophore-conjugated secondary antibody (1:2000, Invitrogen) for 1 h at 37 °C, followed by washing three times with 0.5% TBST. Finally, samples were mounted using ProLong™ Diamond Antifade Mountant with DAPI (Cat# P36970, Invitrogen).

#### Free Fatty Acid (FFA) and Triglyceride (TG) Assays

Free fatty acid and triglyceride levels were quantified using the Free Fatty Acids (NEFA/FFA) Fluorometric Assay Kit (Cat# E-BC-F039, Elabscience®) and the Triglyceride (TG) Fluorometric Assay Kit (Cat# E-BC-F033, Elabscience®), respectively. For each experimental group, 100 oocytes were lysed and processed according to the manufacturer’s protocols. Standards were prepared as instructed to generate calibration curves. Samples and standards were loaded in duplicates into a 96-well OptiPlate, and fluorescence was measured at Ex/Em 535/587–590 nm using an EnSpire Multimode Plate Reader (PerkinElmer) with 500 flashes per well. Raw fluorescence readings were background-corrected against blank wells, and metabolite concentrations were calculated from the standard curves. Final values were normalized to oocyte number and expressed as pmol per oocyte.

#### Mass Spectrometry Sample Preparation

For untargeted whole-cell proteomics, 180–200 oocytes were collected per group (control and DGAT1i-treated). Oocytes were lysed in 2% sodium Deoxycholate (SDC) prepared in 50 mM ammonium bicarbonate (AMBIC) buffer and incubated at 95 °C for 5 min to inactivate endogenous phosphatase and proteases. Lysates were sonicated, centrifuged, and protein concentrations were determined by Bradford assay. Equal amounts of protein were processed for downstream analysis.

Protein extract reduced with 200mM dithiothreitol (DTT) for 1hr at 57 °C at 700RPM and alkylated with 200mM iodoacetamide (IAA) at room temperature in dark for 1 hour to prevent disulphide bond reformation. The samples were then digested using a combination of trypsin and LysC enzymes at 37°C for 16 h. To remove sodium deoxycholate (SDC) from the trypsin-digested samples, 20% formic acid was added to induce precipitation. The mixture was subsequently passed through a 10 kDa filter to eliminate undigested proteins. Sample purification was further performed by desalting with a C18 spin column, following the manufacturer’s protocol. The eluates were then vacuum-dried and reconstituted in 0.3% formic acid. Finally, 1 μg of peptide was injected into the mass spectrometer for analysis. All experiments were conducted with three independent biological replicates.

#### Mass Spectroscopy Data Acquisition

The proteome was analyzed by using the UltiMate 3000 RSLCnano system coupled with the high-resolution Q Exactive HF mass spectrometer (ThermoFisher Scientific). In brief, the full MS scans were carried out by using a resolution value of 60,000, AGC target value of 1 × 106 and acquisition range of 375–1600 m/z with a maximum injection time of 60 ms. The top 25 precursors were selected for the fragmentation. The MS2 acquisition was performed through a resolution of 15,000, AGC target value of 1 × 105, a maximum injection time of 100 ms, an isolation window of 1.3 m/z, and a fixed first mass at 100 m/z. A nonlinear gradient (flow rate of 0.300 μL/min for 180 min) of solvents using 5% of 80% acetonitrile/0.1% formic acid as solvent B and 95% of 0.1% formic acid as solvent A was preferred for eluting the peptides.

#### Mass Spectroscopy Data Analysis

Peptide and protein identification were performed using Proteome Discoverer software (v2.5, Thermo Fisher Scientific, San José, CA, USA) against the UniProtKB/Swiss-Prot *Mus musculus* database (UP000000589).

#### Parameters for Peptide and Protein Search

We selected the dynamic and static modifications of oxidized methionine residues and carbamidomethylation of cysteine residues as our prime search parameters. In addition, we considered the following values of 2, 144, and 6 for most missed cleavage, maximum, and minimum peptide length, respectively. Proteins and peptides with < 1% and < 5% FDR confidence were filtered. The identification of proteins and peptides were based on the 10 ppm and 0.02 Da fragment and precursor mass tolerances. The “Minora Feature Detector,” “Precursor Ions Quantifier,” and “Feature Mapper” nodes workflow were used for the Label-free quantification (based on peptide signals) to identify proteins or peptides (Matthiesen and Jensen <u>2008</u>).

#### Proteomic Data Processing and Analysis

Label-free quantification was performed to determine relative protein abundances across samples. Protein intensity values were normalized by log2 transformation prior to downstream analysis. Venn diagram analysis was used to assess inter-sample variation, while multivariate approaches, including principal component analysis (PCA) and partial least squares discriminant analysis (PLS-DA), were applied to evaluate sample clustering and group discrimination. Multivariate analyses were performed using MetaboAnalyst 6.0 (https://www.metaboanalyst.ca). volcano plot was prepared using prism software. Gene ontology (GO) comparative analysis was performed with Funrich v3.1.3.

#### Transmission Electron Microscopy (TEM) Sample Preparation and Imaging

Goat oocytes (control and DGAT1i-treated) were denuded and fixed with 4% paraformaldehyde (PFA) for 15 min. The oocytes were then embedded in low-melting agarose to facilitate handling. Agarose-embedded oocytes were subjected to primary fixation in 4% PFA and 2.5% glutaraldehyde prepared in 0.1 M sodium cacodylate buffer at 4 °C overnight.

The next day, the fixative was removed, and agarose blocks were washed three times with 0.1 M sodium cacodylate buffer. Secondary fixation was carried out with 1% osmium tetroxide in 0.1 M sodium cacodylate buffer at 4 °C for 1–2 h until the agarose turned black. Afterward, samples were washed three times with 0.1 M sodium cacodylate buffer.

Dehydration was performed through a graded ethanol series: 30%, 50%, 70%, 90%, and 100% (ice-cold), for 30 min each, followed by an additional 100% ethanol wash at room temperature for 1 h. Samples were then infiltrated with epoxy resin: ethanol mixtures in increasing concentrations (1:3 overnight at room temperature, followed by 1:1 for 4–6 h, and then pure epoxy resin for 3–4 h). Finally, the samples were embedded in fresh epoxy resin and polymerized at 65–70 °C for 8 h.

Polymerized resin blocks were trimmed and sectioned using a Leica EM UC7 ultra microtome to obtain ultrathin sections (∼50 nm). Sections were collected on copper grids and contrasted sequentially with UranyLess (10 min; Electron Microscopy Sciences, Cat. No. 22409) and lead citrate (5 min; Electron Microscopy Sciences, Cat. No. 22410). After thorough washing and drying, grids were imaged using a JEM-1400 Flash transmission electron microscope (JEOL, Japan) operated at 80–120 kV.

#### Imaging and quantification

Fluorescence intensity and organelle quantification were performed using Zen Blue software (Zeiss, Germany). Z-stack images were acquired through the entire oocyte at consistent step sizes and acquisition settings; maximum-intensity projections (or individual optical sections where appropriate) were generated for analysis. For intensity and area measurements, individual oocytes were manually outlined with the Graphics Tools → Circle or Spline Contour ROI functions, and mean fluorescence intensity and ROI area were recorded for each oocyte. Subcellular structures — including lipid droplets, endoplasmic reticulum (ER), lysosomes, mitochondria, and PANK2 puncta — were quantified by manual counting using the Point tool, with all images acquired under identical settings. For manual co-localization analysis, dual-stained images were examined, and overlapping structures were identified and marked using the Point tool. All quantification and co-localization analyses were performed blinded to experimental group.

## Statistical analysis

All statistical analyses were performed using GraphPad Prism (version 10.1.2, GraphPad Software, San Diego, CA, USA). Data are expressed as mean ± standard error of the mean (SEM), unless otherwise indicated. Each experiment was performed with two to three independent biological replicates, with the number of oocytes per condition reported in figure legends.

Comparisons between two groups were analysed using unpaired two-tailed t-tests. For comparisons among more than two groups, ordinary one-way ANOVA was performed, followed by either Šídák’s or Tukey’s multiple comparisons test as appropriate. For experiments involving two independent variables two-way ANOVA with Šídák’s multiple comparisons test was applied.

A p-value < 0.05 was considered statistically significant. The specific statistical test applied for each dataset is indicated in the corresponding figure legends.

## Methods contact

Further information and requests for resources and reagents should be directed to the lead contact, H.B.D. Prasada Rao

## ACKNOWLEDGMENTS

We thank the NIAB core Microscope and small animal facility. P.B. was supported by UGC JRF. S.Y. was supported by DBT. A.K. was supported by DBT SRF. A.M. was supported by CSIR SRF. M.A. was supported by UGC SRF. A.K. was supported by UGC SRF. K.K. was supported by ICMR. A.K. was supported by UGC JRF. A. was supported by UGC JRF. S.S was supported by SERB. This work was supported by NIAB core grant C0031 and ICMR grant DDR-IIPRSG-24-RH-15 awarded to H.B.D.P.R.

## AUTHOR CONTRIBUTIONS

P.B., and H.B.D.P.R. conceived the study and designed the experiments. P.B., S.Y., A.K., A.M., M.A., A.K., K.K, A.K., A., S.S., and H.B.D.P.R. performed the experiments and analyzed the data. H.B.D.P.R. and P.B. wrote the manuscript with inputs and edits from all authors.

---

**Model:**
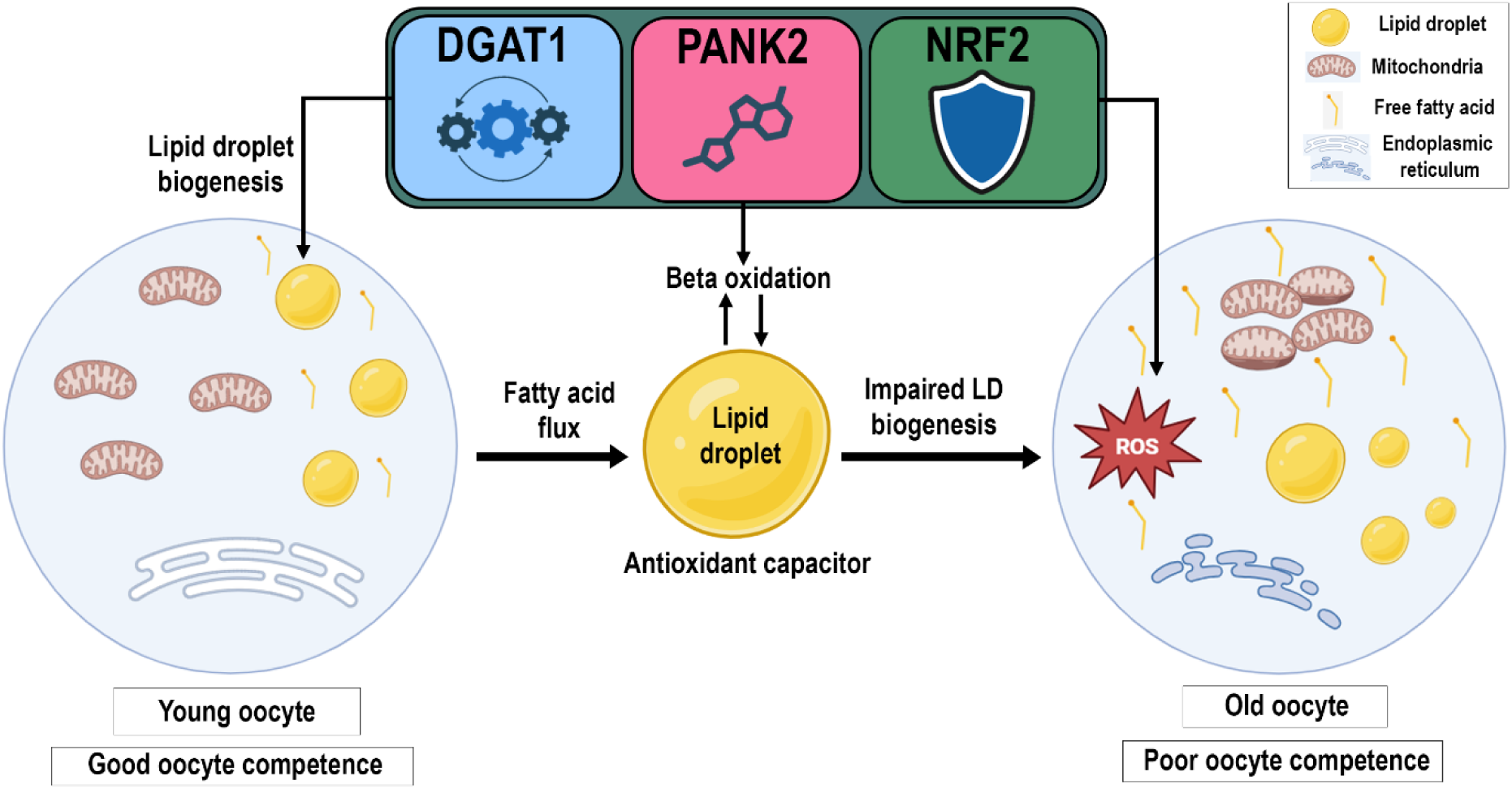
Lipid droplets function as metabolic capacitors in oocytes through the DGAT1–PANK2–NRF2 axis. In young oocytes (left), DGAT1 drives triglyceride synthesis and lipid droplet (LD) biogenesis. LDs act as metabolic capacitors, buffering fatty acid (FA) flux to mitochondria, thereby ensuring controlled β-oxidation and sustaining antioxidant defense through NRF2. This coupling maintains mitochondrial integrity and supports meiotic competence. In contrast, aged or DGAT1-inhibited oocytes (right) exhibit failed LD remodeling and excess FA spillover into mitochondria, resulting in elevated β-oxidation and ROS accumulation. The antioxidant capacity of NRF2 is overwhelmed, leading to mitochondrial dysfunction and meiotic arrest. Together, these findings highlight LDs as dynamic regulators of redox homeostasis and oocyte quality. The figure was made using BioRender.

**Supplementary Fig 1.**
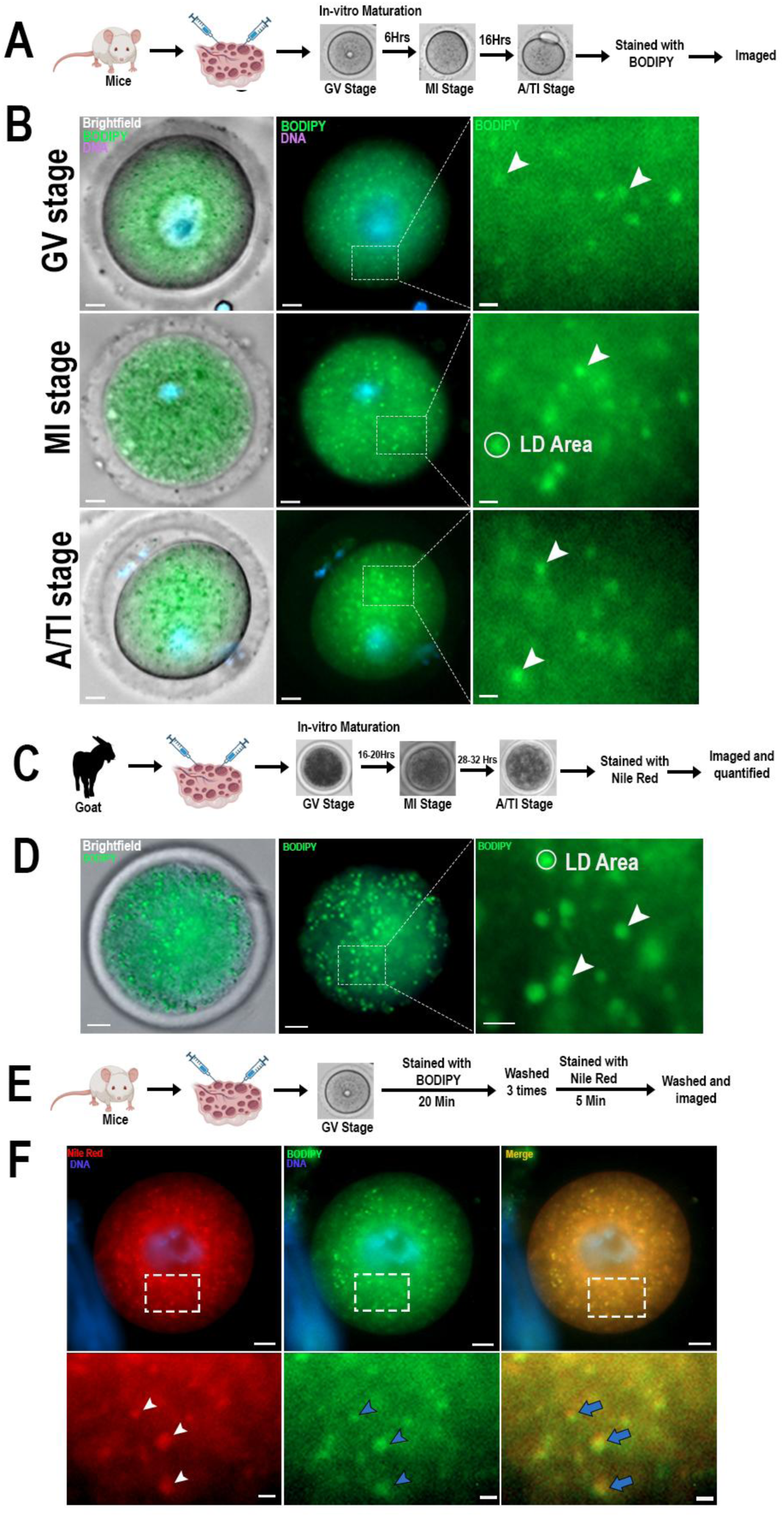
(A) Schematic workflow of mouse oocyte in-vitro maturation (IVM) followed by BODIPY staining. (B) Representative brightfield and fluorescence images of GV-, MI-, and A/TI-stage mouse oocytes stained with BODIPY. White circle indicates LD analysis area; arrowheads mark lipid droplets. (C) Workflow for goat oocyte IVM followed by Nile Red staining. (D) Representative images of GV-stage goat oocytes stained with BODIPY. White circle indicates LD analysis area; arrowheads mark lipid droplets. (E) Workflow of dual staining in GV-stage mouse oocytes with BODIPY and Nile Red. (F) Representative fluorescence images showing BODIPY (green) and Nile Red (red) staining and merged signals. Boxed areas are shown as zoomed regions: white arrowheads indicate LDs (Nile Red), blue arrowheads indicate LDs (BODIPY), and blue arrows indicate colocalization. Scale bars:10 µm (zoomed 2 µm).

**Supplementary Fig 2.**
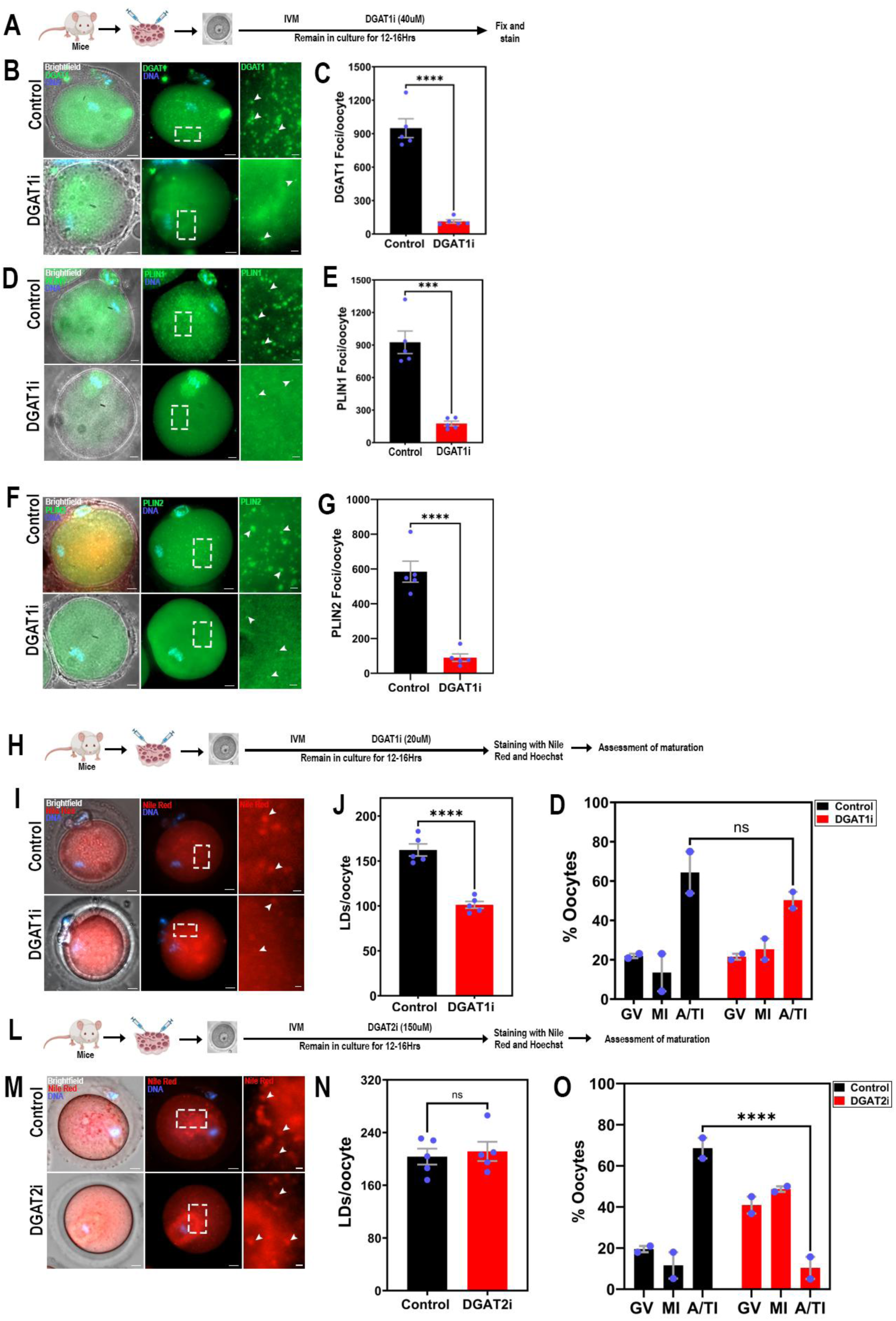
(A) Schematic workflow for DGAT1 inhibitor (DGAT1i) treatment during in vitro maturation (IVM) followed by immunostaining. **(B)** Representative images of control and DGAT1i-treated oocytes stained for DGAT1 (green) and Hoechst (DNA, blue). Boxed regions show zoomed areas with DGAT1 puncta (arrowheads). **(C)** Quantification of DGAT1 foci per oocyte. **(D)** Representative images of control and DGAT1i-treated oocytes stained for PLIN1 (green). Boxed regions show zoomed areas with PLIN1 puncta (arrowheads). **(E)** Quantification of PLIN1 foci per oocyte. **(F)** Representative images of control and DGAT1i-treated oocytes stained for PLIN2 (green). Boxed regions show zoomed areas with PLIN2 puncta (arrowheads). **(G)** Quantification of PLIN2 foci per oocyte. **(H)** Schematic workflow for DGAT1i treatment during IVM followed by Nile Red staining and maturation assessment. **(I)** Representative brightfield and fluorescence images of control and DGAT1i-treated oocytes stained with Nile Red (LDs, red). Boxed regions show zoomed LDs (arrowheads). **(J)** Quantification of LD number per oocyte. **(K)** Assessment of oocyte maturation shown as percent oocytes at GV, MI, and A/TI stages after DGAT1i treatment. (L) Schematic workflow for DGAT2i treatment during IVM. **(M)** Representative images of control and DGAT2i-treated oocytes stained with Nile Red (red). Boxed regions show zoomed LDs (arrowheads). **(N)** Quantification of LD number per oocyte. **(O)** Assessment of oocyte maturation shown as percent oocytes at GV, MI, and A/TI stages after DGAT2i treatment. Maturation experiments were performed in duplicate with 20–30 oocytes per group per replicate. Data are presented as mean ± SEM. Statistical significance was assessed using unpaired t test or two-way ANOVA with Šídák’s multiple comparisons test. Statistical significance is indicated as follows: ***p < 0.001; ****p < 0.0001; ns = not significant. Scale bars:10 µm (zoomed 2 µm).

**Supplementary Fig 3.**
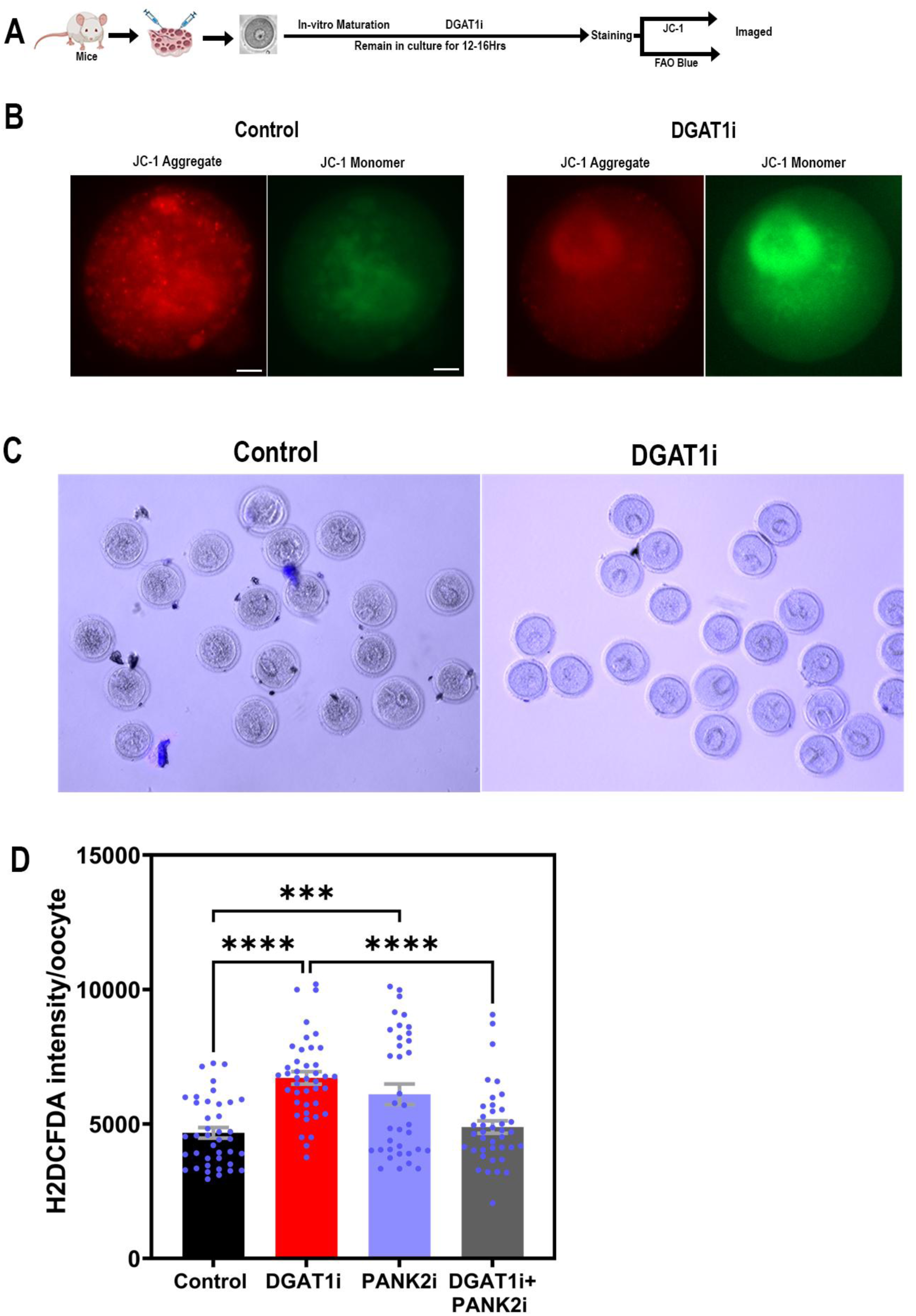
(A) Schematic workflow of DGAT1 inhibitor (DGAT1i) treatment during in vitro maturation (IVM) followed by JC-1 and FAO Blue staining. **(B)** Representative fluorescence images of control and DGAT1i-treated oocytes stained with JC-1 showing mitochondrial aggregates (red) and monomers (green). **(C)** Representative FAO Blue staining images showing fatty acid β-oxidation activity in control and DGAT1i-treated oocytes. **(D)** Quantification of ROS levels using H2DCFDA intensity per oocyte in control, DGAT1i, PANK2i, and DGAT1i+PANK2i groups. Data are presented as mean ± SEM. Statistical significance was assessed using ordinary one-way ANOVA with Šídák’s multiple comparisons test. Statistical significance is indicated as follows: ***p < 0.001; ****p < 0.0001; ns = not significant. Scale bars:10 µm.

**Supplementary Fig 4.**
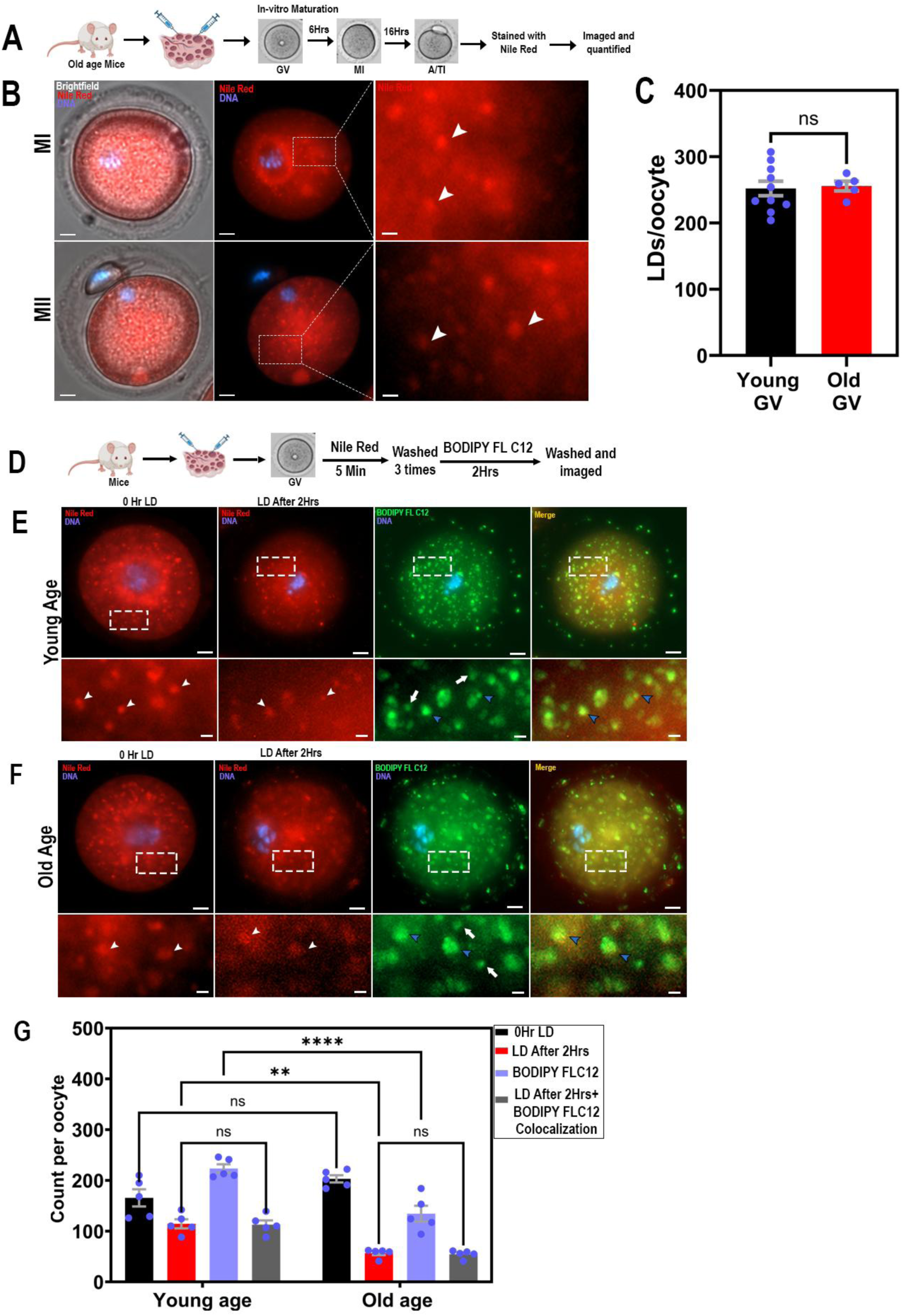
(A) Schematic workflow for assessing lipid droplets (LDs) during in vitro maturation (IVM) in old mice using Nile Red staining. **(B)** Representative brightfield and fluorescence images of MI- and MII-stage old oocytes stained with Nile Red. Boxed areas are shown as zoomed images with white arrowheads indicating LDs. (C) Quantification of LD number per GV-stage oocyte in young and old C57BL/6J (Black6J) mice. **(D)** Schematic workflow for dual staining of GV oocytes with Nile Red and BODIPY FL C12. **(E–F)** Representative fluorescence images of GV-stage oocytes from young (E) and old (F) C57BL/6J (BL6) mice stained with Nile Red (red, LDs) and BODIPY FL C12 (green, de novo LD synthesis). Each group shows four panels: (1) Nile Red at 0 hr (pre-existing LDs); (2) Nile Red after 2 hr; (3) BODIPY FL C12 after 2 hr; (4) merged image showing colocalization. Boxed areas are shown as zoomed images: white arrowheads indicate LDs, blue arrowheads indicate BODIPY FL C12 colocalized with Nile Red, and white arrows indicate newly synthesized LDs. **(G)** Quantification of LDs (Nile Red at 0 hr), LDs after 2 hr Nile Red staining, newly synthesized LDs (BODIPY FL C12 after 2 hr), and colocalized LDs in GV-stage oocytes from young and old C57BL/6J (BL6) mice. Data are presented as mean ± SEM. Statistical significance was assessed using an unpaired t test or two-way ANOVA with Šídák’s multiple comparisons test. Statistical significance is indicated as follows: ***p < 0.001; ****p < 0.0001; ns = not significant. Scale bars:10 µm (zoomed 2 µm).

